# Demographic responses to climatic changes during the Final Palaeolithic in Europe

**DOI:** 10.1101/2024.09.11.612499

**Authors:** Isabell Schmidt, Birgit Gehlen, Katja Winkler, Alvaro Arrizabalaga, Nico Arts, Nuno Bicho, Philippe Crombé, Berit Valentin Eriksen, Sonja B. Grimm, Katarina Kapustka, Mathieu Langlais, Ludovic Mevel, Nicolas Naudinot, Zdeňka Nerudová, Marcel Niekus, Marco Peresani, Felix Riede, Florian Sauer, Werner Schön, Iwona Sobkowiak-Tabaka, Hans Vandendriessche, Mara-Julia Weber, Annabell Zander, Andreas Zimmermann, Andreas Maier

## Abstract

The European Final Palaeolithic witnessed marked changes in almost all societal domains. Despite a rich body of evidence, our knowledge of palaeodemographic processes and regional population dynamics still needs to be improved. In this study, we present regionally differentiated estimates of absolute numbers and population densities for the Greenland Interstadial 1d-a (GI-1d-a; 14-12.7 ka BP) and the Greenland Stadial 1 (GS-1; 12.7-11.6 ka BP) for Southern, Western, Northern and Central Europe. The data were obtained by applying the Cologne Protocol, a geostatistical approach for estimating prehistoric population size and density, to a newly compiled dataset of Final Palaeolithic sites. On a large spatio-temporal scale, we observe a shift of the main areas of human occupation from the Franco-Cantabrian region, which was intensely occupied during most phases of the preceding Upper Palaeolithic, to regions north of the Alps. At smaller scales, we observe divergent regional trends in the Final Palaeolithic meta-population: during GI 1d-a, a decreasing population in southwestern Europe and an increasing population in north-eastern Central Europe. For the first time since the dispersal of anatomically modern humans into Europe, we see that Central Europe becomes the dominant demographic growth area. Subsequently, the climatic cooling of GS-1 coincides with a pronounced population decline in most parts of the study area. An apparent increase in population density occurs only in north-eastern Central Europe and north-eastern Italy. Our estimates suggest that the total population was reduced by half. Similar results, with a relationship between decreasing temperatures and decreasing populations, have already been observed for the late phase of the Gravettian, when populations were reduced to only one third of those estimated for the early phase. Yet, in contrast to the collapse of local populations during the late Gravettian, the increase in population densities in Central Europe during GS-1 indicates population movements eastwards, possibly in response to deteriorating climatic conditions, particularly in western regions during the Younger Dryas.

## Introduction

The Late Glacial, comprising Greenland Interstadial 1 (GI-1) and Stadial 1 (GS-1), was a period when the impact of rapidly changing environmental conditions drove high-magnitude and high-amplitude ecosystem shifts. The extensive dispersal of shrub and tree species into northern Western and Central Europe alongside the associated fauna during GI-1 contributed to creating a European habitat with particularly pronounced regional differences. Although the so-called ‘mammoth steppe’ [1] of preceding periods was also never a uniform habitat [2], this emerging, regionally diverse ecotonal patchwork presumably had consequences for most fundamental properties of hunter-gatherer societies, such as subsistence strategies, social networks, and – prominently – the structure and distribution of the underlying meta-population.

In Europe, these changes are culturally associated with the transition from the Late Upper to the Final Palaeolithic period (e.g., [3–6]). Subsistence strategies change from hunting migratory game to a preference for more stationary, thermophilous game. Settlement and land use patterns are marked by sites with functionally differentiated areas, such as Le Closeau or Reichwalde [7,8] rather than large agglomerative sites, such as Gönnersdorf [9–11]. Changes in lithic technology, particularly in core reduction strategies, signalled a less normative use of lithic raw material of varying quality, alongside a regionalisation in lithic projectile morphology [3,12–18]. Visual representations are no longer found in caves but begin to appear frequently on pebbles. In addition, abstract representations become much more dominant than figurative ones. From a demographic perspective, these profound changes went hand in hand with expanding hunter-gatherer groups into the northern European lowlands. From the extensive dispersal of anatomically modern humans into Europe during the Aurignacian up to the repopulation of Central Europe after the Last Glacial Maximum, the meta-population in Europe had undergone substantial fluctuations regarding absolute numbers and densities of people as well as social interconnectedness of sub-populations and spatial distributions of genetic haplotypes [19–23]. In its broadest outline, however, the large-scale distribution pattern of hunter-gatherers remained remarkably stable, with one demographic centre in the Franco-Cantabrian region and another in eastern Central Europe, covering Lower Austria, Moravia, southern Poland, and northern Hungary. In the Final Palaeolithic, this pattern changed drastically.

This change occurred between 15 and 14 ka BP, but there is no generally accepted date or event marking the end of the Late Upper and the beginning of the Final Palaeolithic in Western and Central Europe. Particularly problematic for a chronological delimitation is a plateau in the calibration curve during this period that causes wide calibration age ranges [24,25] and, thus, reduces chronological resolution. Further complicating matters are regionally differentiated environmental responses to the climatic shift at the transition from the Late Pleniglacial to GI-1e, which appears rather abrupt in the Greenland ice core records and may have been related to a substantial meltwater discharge into the North Atlantic [26] and tipping points in sea-ice cover [27]. Atmospheric warming over Europe would have started well before, and the associated ecosystem change likely was more gradual [28,29]. There is growing evidence that such an early start and comparatively slow change of ecosystems did not only take place south of the Alps [30,31], but also in Central Europe [32,33]. Variable response periods of the vegetation in diverse parts of Europe (e.g., [33–36]) further indicate regionally distinct ecosystem reactions differentiated by both leads and lags. Such differences are not limited to warming events but also occurred during the cooling of GS-1 (≈ Younger Dryas) and subsequent warming when de-and increases in seasonal temperature and precipitation varied markedly between different areas throughout Europe [37–41]. An earlier and more gradual ecosystem change prior to GI-1e also has consequences for our understanding of the timing of human population dispersal into Northern Europe. It is in accord with an apparently earlier northward expansion of hunter-gatherers of the Creswellian and Hamburgian [42–44].

From a palaeogenetic point of view, the limited data of this period indicates a major turnover between 15 and 14 ka BP, during which the so-called ‘Oberkassel cluster’ became the dominant haplogroup in Central Europe [23]. Thus, while it is difficult to pinpoint the start of the Final Palaeolithic, it is undisputed that archaeological phenomena after 14 ka BP (post-GI-1e) are attributed to the Final Palaeolithic [3,18,45,46]. Therefore, we chose this date as the older boundary of our study, while the younger boundary coincides with the end of GS-1 and the end of the Pleistocene at 11.6 ka BP.

Here, we present regionally differentiated estimates of absolute numbers and densities of people for two phases of the Final Palaeolithic according to the Cologne Protocol [47]. A previous study considered the Final Palaeolithic as a whole [21], but limited data granularity has hitherto precluded analyses at a finer resolution. Through the involvement of regional experts from different countries, we compiled an extensive database [48], allowing for a chronological subdivision of the assemblages into two phases. The older phase, dates between 14 and 12.7 ka BP and thus corresponds to GI-1d-a or the Older Dryas and the Allerød *sensu* Iversen [49], while the younger phase dates to between 12.7 and 11.6 ka BP and thus corresponds to GS-1 or the Younger Dryas (**Fig 1**). Based on our estimates, we address how the human meta-population responded demographically to environmental change associated with the climatic cooling from GI-1d-a to GS-1 at a subcontinental and regional spatial scale. We compare these results with the demographic response during the Gravettian [50], that was similarly linked to major cooling, and discuss similarities and differences, as well as implications for social networks and behavioural adjustments.

**Fig 1.**
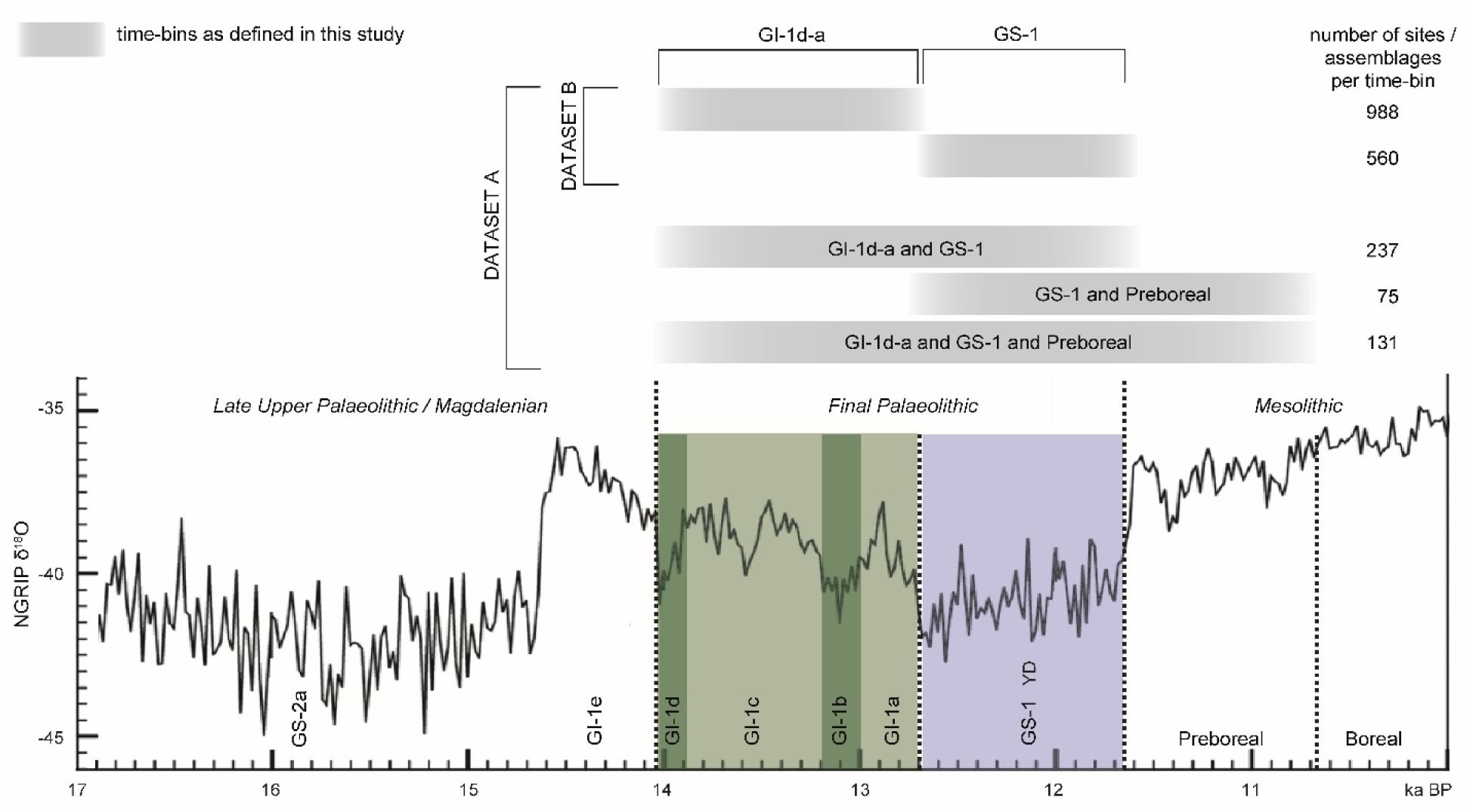
Chronological framework of this study. The time-bins defined in the upper part of the graph indicate the different typo-chronological resolutions of the assemblages that were taken into account for the database and analysis.

## Material and Methods

The database underlying this study comprises archaeological sites from Southern, Western, Northern and Central Europe. It has been compiled with the regional expertise of the participating authors and is available online as an open-access resource [48]. To benchmark this detailed recording of the evidence across the study area, unpublished (and thus not verifiable) material and single finds were not considered. Some differences, however, turned out to be inevitable due to data accessibility (publication, database management of heritage authorities) or regionally varying properties of the record (e.g., chronological resolution). In sum, the database considers material from 1788 sites (**Fig 2**, excavations and surface collections). 1548 assemblages can either be directly dated and/or typologically or stratigraphically assigned to one of the two phases. GI-1d-a (14-12.7 ka BP) comprises 988 assemblages and GS-1 (12.7-11.6 ka BP), comprises 560 assemblages. Material from assemblages that cover larger time-bins (see **Fig 1**) contain 237 assemblages assigned to the Final Palaeolithic in general, 131 assemblages assigned to the Final Palaeolithic and the Preboreal in general, and 75 assemblages are recorded separately because their chronological placement relates to the final GS-1 and the succeeding Preboreal phase. Frequent generic attribution is most problematic for evidence from Central Europe.

**Fig 2.**
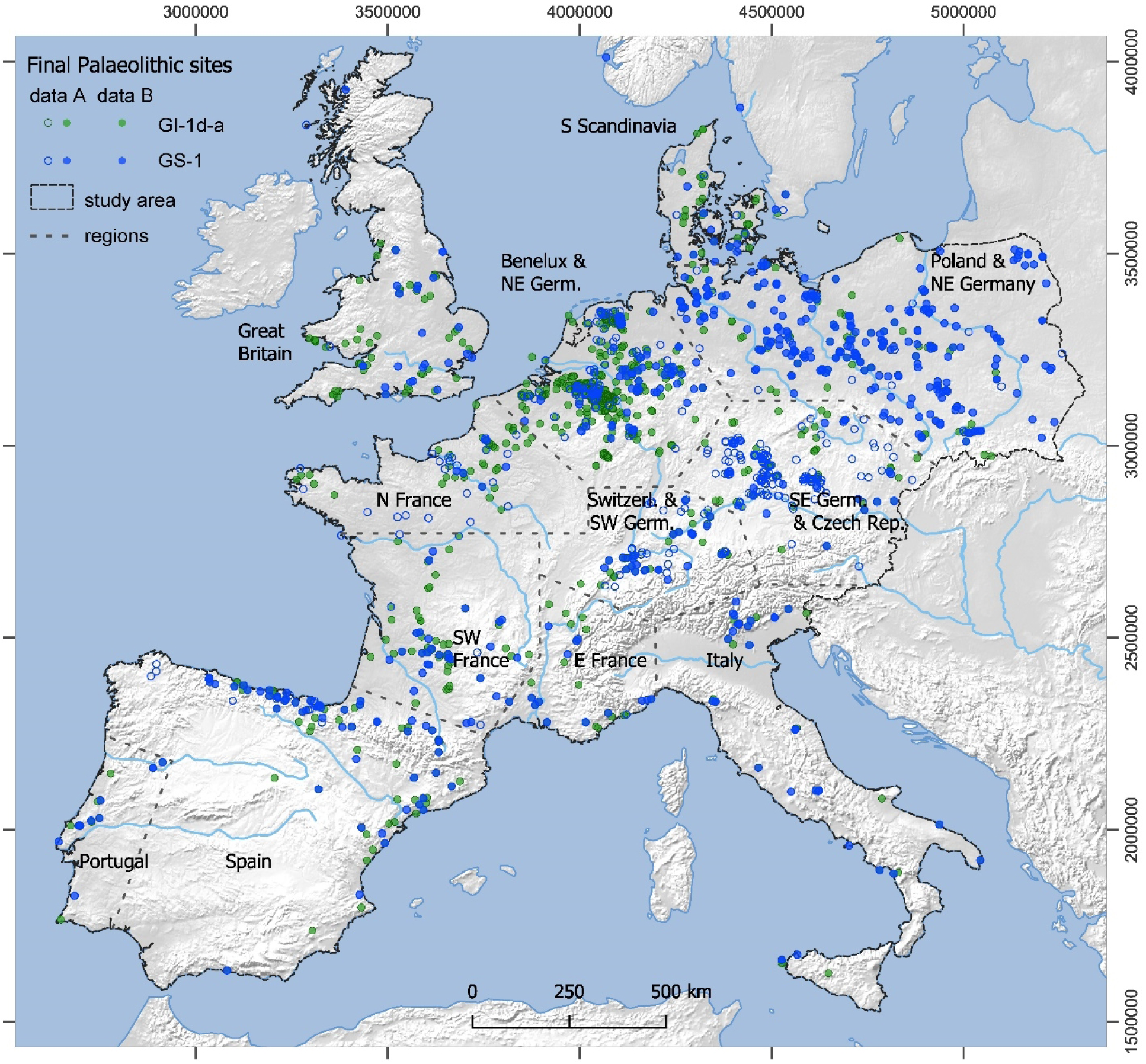
Distribution of Final Palaeolithic sites in the study area. Green dots: GI-1d-a; blue dots: GS-1. Filled circles: dataset B (chronologically subdivided sites); filled and open circles: dataset A (all sites; see text and Fig 1. Base map details: see supporting information).

Submerged coastal areas and so-called Doggerland provided large, formerly inhabitable landmasses. Evidence of human occupation from these areas is extremely scarce and, therefore, virtually absent in the dataset. Examples of long-distance raw material transport, e.g., of Helgoland flint [51–53], speak to human presence across these now submerged areas. If diagnostic archaeological artefacts are recovered from below current sea level, they often lack geospatial information. Our calculations of the estimates, regional comparisons and subsequent interpretations of the results therefore rest upon the available evidence from the current landmass, as mapped in **Fig 2**.

On the North European Plain and adjacent areas, GI-1d-a mainly comprises sites attributed to the *Federmesser-Gruppen* or *Federmesser* culture (e.g., [54–59]). The beginning of this phase overlaps slightly with the Havelte phase of the Hamburgian [42,46,56,60,61] and potentially with the latest Creswellian sites [54,62,63]. Given that most sites attributed to these industries predate GI-1d-a (see also [64]), Havelte and Creswellian assemblages were consistently excluded from the dataset. The disappearance of these ‘cultures’ may in itself be indicative of a demographic break at this important tipping-point transition [44,56,65] as indicated by results from the British Isles [66]. During GS-1, assemblages assigned to the Ahrensburgian and Swiderian occur in the Central and Northern lowlands [58,67–69], at time persisting into the subsequent Preboreal period (e.g., [70]). The position of the Bromme culture in this framework, situated at the transition from Allerød to Younger Dryas, is subject to discussion [56,71,72]. A high number of surface collections characterises the archaeological evidence on the northern Plains in general.

In the Benelux, Ahrensburgian and Epi-Ahrensburgian assemblages [73–77] replace the *Federmesser-Gruppen* during GS-1, with most dates placing the Ahrenbsurgian notably late into GS-1 or even into the Holocene [78]. Long Blade industries [79,80] and Belloisian/Long Blade assemblages [80–84] emerged in the UK and the Paris Basin, respectively. Assemblages with so-called ‘Riesenklingen’ (giant blades) also occur in the Netherlands [76], north-western Germany (e.g., [85]) and Denmark [86], where they are assigned to the younger half of GS-1 and occasionally seem to diffuse into the Preboreal [87].

In France, the term ‘Azilian’ is used in competition with ‘Federmesser-Gruppen’. The latter is more commonly used in northern France, particularly in the Somme valley. This distinction has more to do with taxonomy and the history of research [88] than with any archaeological reality. However, differences cannot be overlooked typologically and technologically, especially when comparing certain sites (e.g., Le Closeau *vs* Saleux), although it is not yet possible to explain the exact meaning of this variability. This taxonomic question also applies to the ‘Belloisian’, ‘Laborian’ and ‘Epi-Ahrensbourgian’ entities, which are sometimes supposed to be part of the same technical trend (‘FBBT’; [89–92]). The situation is, of course, different for assemblages related to the different phases of the Epigravettian period, and particularly to the most recent ones. They reflect the probable spread of these traditions beyond southeastern France -at least along the Rhone corridor and certainly as far as the Jura range [93–96] -during the GS-1 period. In the actual state of knowledge, we can highlight offsets between their chronometric boundaries and typological changes [83,90,97–99].

Along the Atlantic façade of the Iberian Peninsula, regional chronologies differ in the assignment of taxonomic units to certain periods [100–103]. In Portugal, the Magdalenian is seen as ending before the Younger Dryas. The following phase is frequently called Epipalaeolithic [102,104] or Early Mesolithic [103]. Either way, the lithic assemblages appear very similar in the two phases in both technology and typology. In Cantabrian Spain and Pyrenees, Azilian is the most used term.

In Italy, complexes dated to the Late Glacial interstadial and the Younger Dryas are framed within the late and terminal Epigravettian, although differences exist between regions in chronology and terminology [105,106]. Indeed, from the end of the Last Glacial Maximum up to the Pleistocene/Holocene boundary, the Epigravettian modified its technological features, the typology of tools and of weapons, as related to the use of bow and arrow, particularly in the latest phases [107]. The shortening of certain tools, such as frontal end-scrapers [108], and the spread of backed knives occur alongside with the appearance of geometric microliths and the increasing use of the microburin blow technique. These modifications seem to be shared at larger scale in Eastern and Mediterranean Europe [12].

The greatest difficulties for a chronological subdivision were encountered along the northern range of the Alps, particularly in southern Germany and the Czech Republic. In contrast to many other areas, there is no discernible shift in typology between GI-1d-a and GS-1 in south-eastern Germany, and assemblages of both phases are attributed to the *Federmesser-Gruppen* [109]. The absence of chronometric data further complicates matters since most assemblages are surface collections and organic preservation is poor. In the Czech Republic, *Federmesser-Gruppen* are only present in the north-western part of Bohemia towards the end of GS-1 [110,111]. In contrast, assemblages of GI-1d-a and the early GS-1 and in other parts of the country are at times assigned to the Epimagdalenian or Final Palaeolithic. However, many sites are surface collections and can therefore not be securely associated with either of the two phases [112,113]. In the sandstone regions, stratified and radiometrically dated sites are known [33,114,115], which may help refine the chronology of the Final Palaeolithic in this region. Better dated sites and assemblages occur in southernmost Germany and Switzerland [116–121].

Alongside the database on Final Palaeolithic sites and assemblages, we collected data on lithic raw material transport from the literature. Where possible, the information was divided into the two chronological phases (**Fig 1**), too. If a site was known to contain assemblages from both periods but raw material data are presented in the literature generically as ‘Final Palaeolithic’, these data were assigned to both periods equally. If a site was known to be only generically assigned to the Final Palaeolithic, the raw material data was recorded but not considered for the phase-specific calculation of the demographic estimates. This accounts especially for sites from the area of SE Germany and the Czech Republic (e.g., [109,110]). We also excluded raw materials documented by single pieces and only considered raw materials representing at least 1% of an assemblage (see also [47]). In a second step, we determined the Raw Material Catchment Areas (RMCA) for each assemblage by adding a 5 km radius buffer around the archaeological site and constructing a convex hull around the site-buffer and the related raw material sources. We excluded RMCAs which only comprised a single source-to-site distance, as well as RMCAs which were smaller than 500 km^2^ since they turned out to be exceptional outliers among the overall RMCA sizes (these assemblages are from Grotte du le Bichon and Blot for GI-1d-a and Gahlen-Schermbeck, Bad Breisig and Babí Pec for GS-1). Eventually, for GI-1d-a, 53 out of 108 RMCAs and for GS-1, 39 out of 97 RMCAs were used to calculate demographic estimates (**S1 and S2 Table**). The scarcity or even lack of available raw material data in some Core Areas (e.g., Great Britain, Denmark, and northern Spain) required transferring RMCA-sizes from adjacent areas.

The calculation of population estimates follows the Cologne Protocol [47]. In brief, it statistically divides a certain map section, called Total Area of Calculation, into areas of high hunter-gatherer activities, called Core Areas, and areas of little or no activities. The Core Areas are delimited by calculating areas with equal site densities, encircled by the so-called optimally describing isoline. To account for differences in site density within Europe, the optimally describing isolines have been modelled individually for three regions (for details, see **S1 and S2 Figs**). The Total Area of Calculation for the current study is 2.6 million km^2^ of Southern, Western, Northern, and Central Europe, following the current coastlines. To estimate the number of hunter-gatherer groups per Core Area, the size of each optimally describing isoline (in km^2^) is divided by the quartiles (Q1, Q2, Q3) of the related RMCA. The results are then multiplied with an average group size (42.5) derived from ethno-historic cases selected based on socio-economic considerations (see [47,122]), providing the estimate of the absolute number of people. Population densities are then calculated for each Core Area by dividing the respective numbers of people by the size of the Core Area, and for the Total Area of Calculation by dividing the number of all people by the size of the selected map section.

To account for the fact that a chronological subdivision of the dataset was more feasible in some regions and less so in others, we decided to calculate for each phase two demographic estimates based on different datasets: the so-called ‘dataset A’ includes all sites, subdivided assemblages as well as sites that are only generically assigned to the Final Palaeolithic period or the time-bins indicated in **Fig 1**. Information about human presence recorded from sites with low chronological resolution is thus retained in the modelling, which may then, however, lead to overestimations for a given phase at regional scales. The second demographic estimate, based on ’dataset B‘, in contrast, considers only assemblages with a clear chronological subdivision assigned to them, attributable either to GI-1d-a or to GS-1, or sites that can be clearly assigned to both phases. Due to the overall smaller sample size of ‘dataset B’, it is expected to produce lower estimates – likely underestimating the number of people – than the ones derived from dataset A, but it provides the best available high chronological control. All estimates are given as a mean estimate and, based on our calculation protocol, with a minima and maxima estimate (related to Q1 and Q3 of the RMCA).

## Results

In the following, we will discuss the results of the estimates from dataset A and dataset B for each chronological phase, first considering dataset A with all sites, and then the smaller but chronologically highly resolved dataset B.

For the <u>estimates from dataset A</u>, all Core Areas of GI-1d-a (**Fig 3**) enclose an area of approximately 635,000 km^2^ and contain 90% of the sites (1220 out of 1356), while those of GS-1 (**Fig 4**), in contrast, enclose only about 481,000 km^2^ and contain 82% of the sites (825 out of 1003; for further details, see **S1 Fig**). The mean estimate for the total population for GI-1d-a amounts to roughly 8100 people at any given time, with minima and maxima ranging from 4800 to 14,300 (**Table 1**). For the GS-1, the mean estimate for the total population drops to 4250 people living at the same time, with minima and maxima ranging from 2200 to 8800 people (**Table 2**). This decrease in population size is accompanied by a decrease in population density within the Core Areas from a mean value of 1.3 people per 100 km² (p/100km²) in GI-1d-a (min. 0.8, max. 2.2) to 0.9 p/100km² in GS-1 (min. 0.5, max. 1.8). In the regional results, the Czech Republic and SE Germany are combined by the Core Area and have, with around 2500 people, the largest and with 2.2 p/100 km² also the densest population in GI-1d-a. However, they also show the strongest decline to only 460 people and 0.9 p/100 km² in GS-1. The second highest estimates for GI-1d-a are found in northern Spain and the French Pyrenees with 1200 people, closely followed by Poland and NE Germany with 1100 people. Despite these similar numbers, the density varies considerably between 1.7 p/100 km² in the former and 0.8 p/100 km² in the latter areas. For the total area of calculation (‘study area’ in **Fig 2**), the estimated population density drops from 0.003 in GI-1d-a to 0.002 p/km^2^ in GS-1.

**Fig 3.**
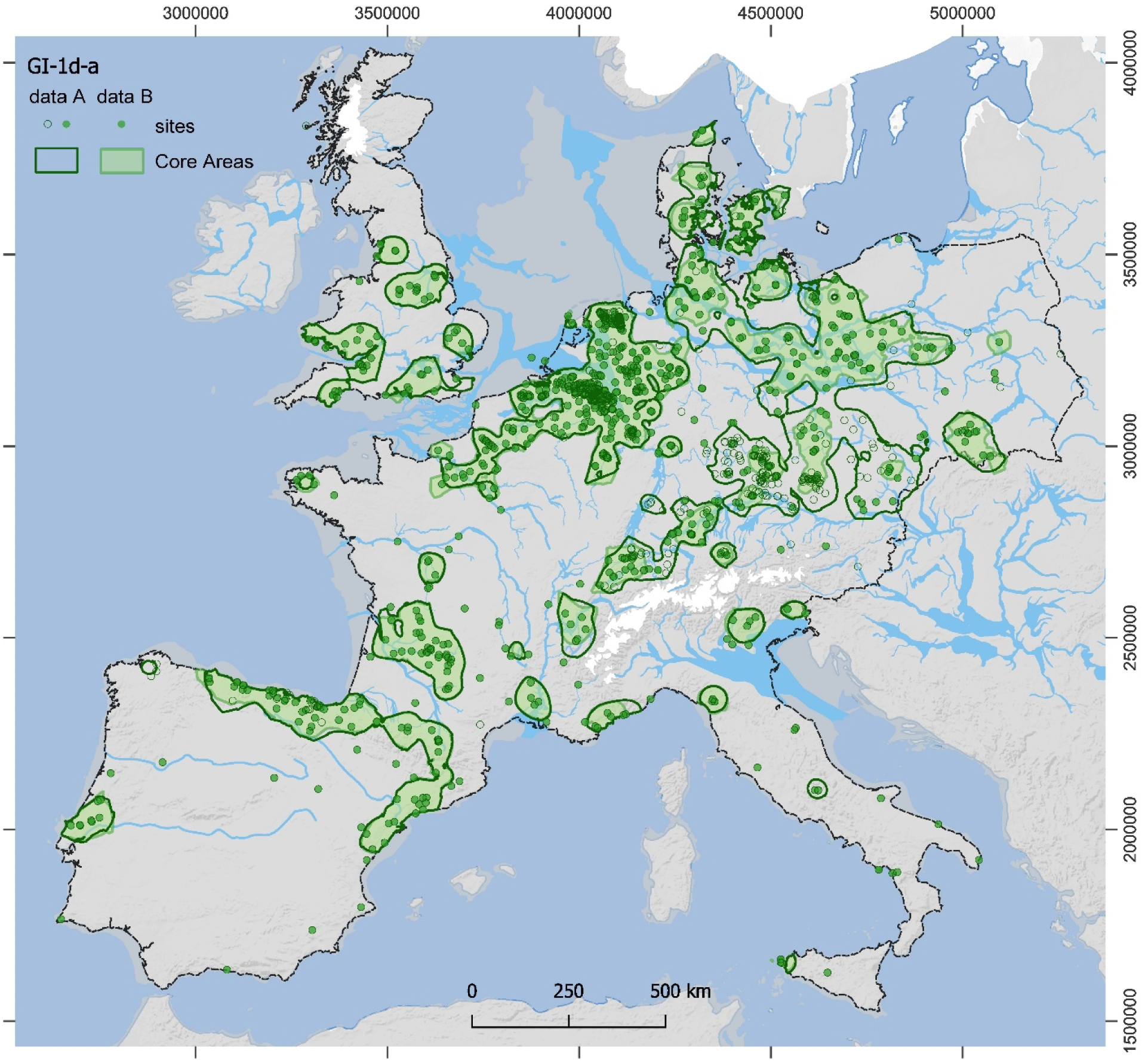
Site distribution and modelled Core Areas for GI-1d-a. The data from dataset A (thick lines, empty and filled dots) and dataset B (shaded areas, filled dots only) are shown. For details on areal data and population estimates see **Table 1** (data A) and **S3 Table** (data B). Reconstructions of ice sheets, river channels, and coastline (>45°N) correspond to GI-1c-a conditions (see supporting information and [3] and references therein).

**Fig 4.**
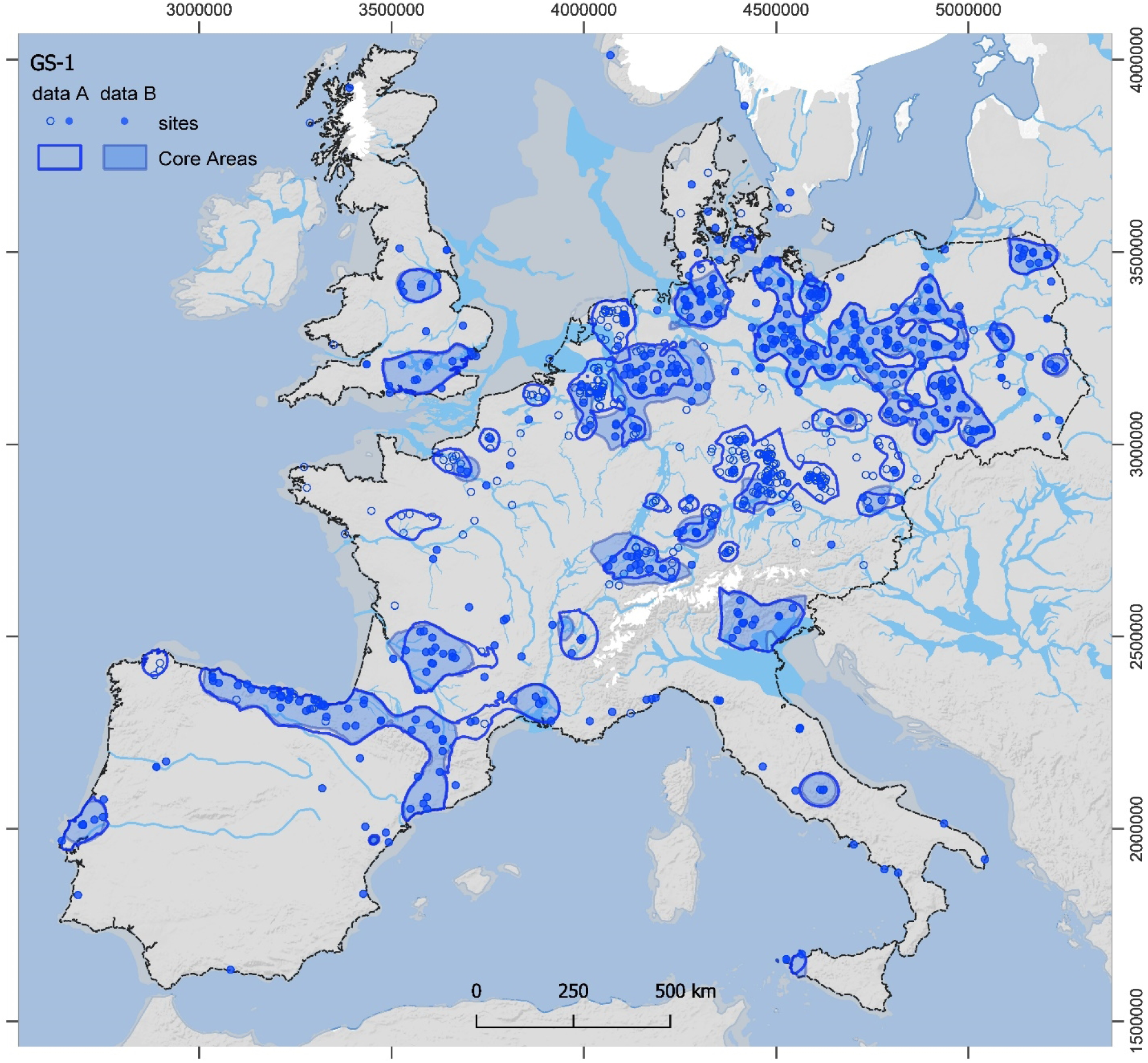
Site distribution and modelled Core Areas for GS-1. The data from dataset A (thick lines, empty and filled dots) and dataset B (shaded areas, filled dots only) are shown. For details on areal data and population estimates see **Table 2** (data A) and **S4 Table** (data B). Reconstructions of ice sheets, river channels, and coastline (>45°N) correspond to GS-1 conditions (see [3] and references therein).

**Table 1.**
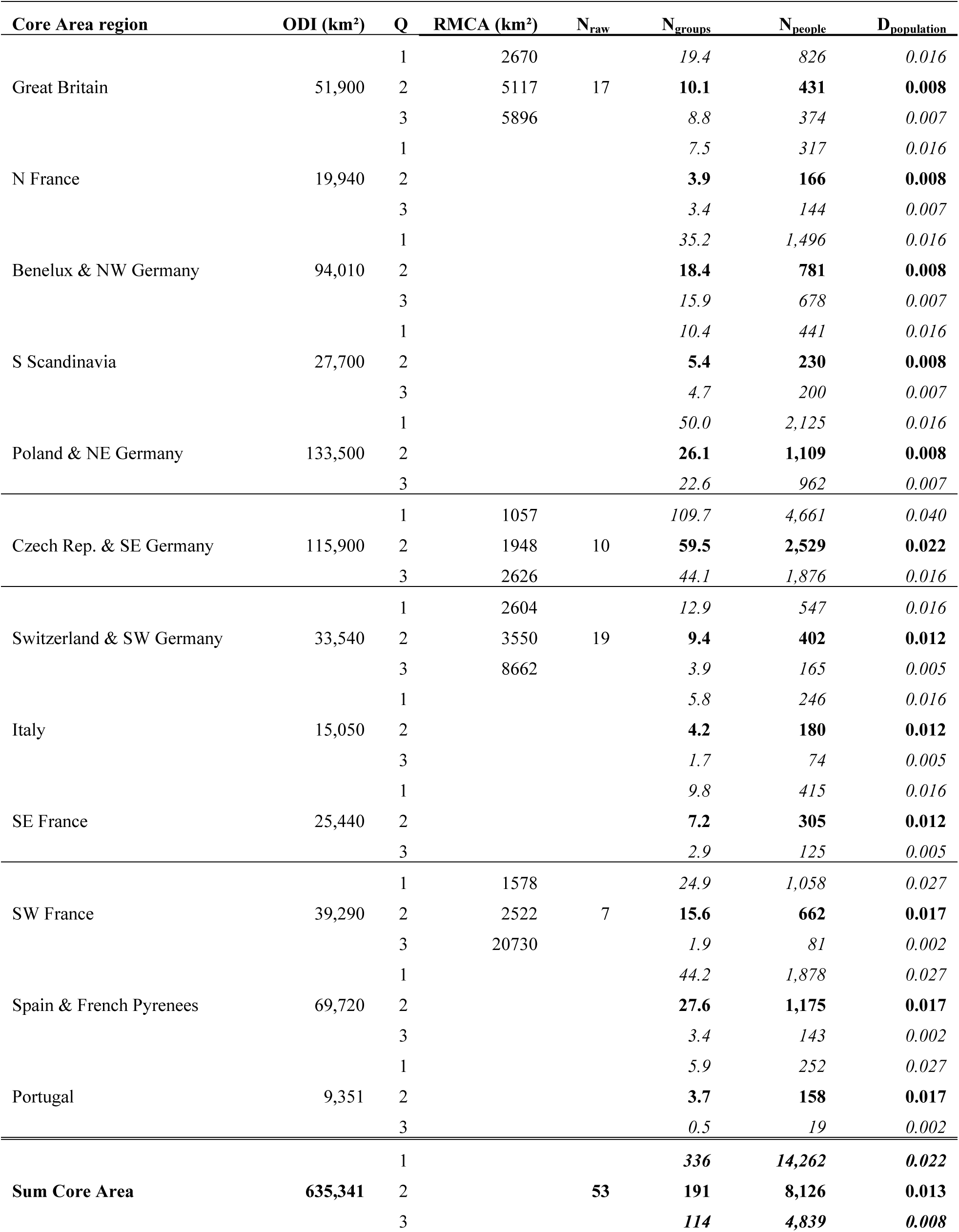
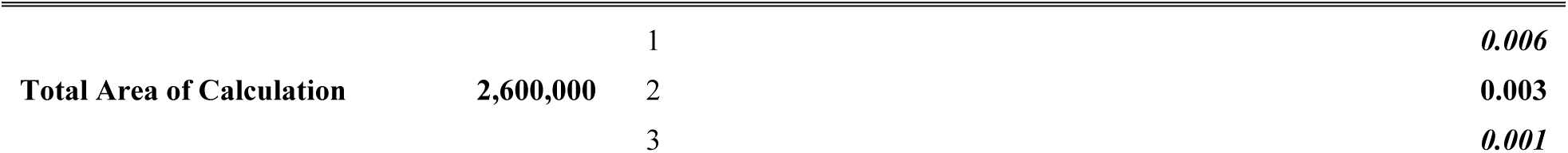
Regionally distinguished demographic estimates for GI-1d-a based on dataset A (all sites). Thin horizontal lines indicate regions with transfer of RMCA data (see **S3 Fig**). ODI = optimally describing isoline in km²; Q = Quartile of RMCA [1 = lower, 2 = median, 3 = upper]; RMCA (km²) = raw material catchment area in km²; N_raw_ = number of RMCAs; N_groups_ = number of groups; N_people_ = number of people; D_population_ = population density within Core Areas as p/km2, and the Total Area of Calculation (bottom rows).

**Table 2.** Regionally distinguished demographic estimates for GS-1 based on dataset A. Thin horizontal lines indicate regions with transfer of RMCA data (see **S4 Fig**). For an explanation on abbreviations see Table 1.

| **Core Area region** | **ODI (km<sup>2</sup>)** | **Q** | **RMCA (km<sup>2</sup>)** | **N<sub>raw</sub>** | **N<sub>groups</sub>** | **N<sub>people</sub>** | **D<sub>population</sub>** |
| --- | --- | --- | --- | --- | --- | --- | --- |
|  |  | 1 | 2934 |  | 8.8 | 374 | 0.014 |
| Great Britain | 25,830 | 2 | 5187 | 4 | **5.0** | **212** | **0.008** |
|  |  | 3 | 7462 |  | 3.5 | 147 | 0.006 |
|  |  | 1 |  |  | 4.8 | 202 | 0.014 |
| N France | 13,950 | 2 |  |  | **2.7** | **114** | **0.008** |
|  |  | 3 |  |  | 1.9 | 79 | 0.006 |
|  |  | 1 |  |  | 17.9 | 761 | 0.014 |
| Benelux & NW Germany | 52,500 | 2 |  |  | **10.1** | **430** | **0.008** |
|  |  | 3 |  |  | 7.0 | 299 | 0.006 |
|  |  | 1 |  |  | 0.5 | 20 | 0.014 |
| S Scandinavia | 1,350 | 2 |  |  | **0.3** | **11** | **0.008** |
|  |  | 3 |  |  | 0.2 | 8 | 0.006 |
|  |  | 1 |  |  | 50.8 | 2,159 | 0.014 |
| Poland & NE Germany | 149,000 | 2 |  |  | **28.7** | **1,221** | **0.008** |
|  |  | 3 |  |  | 20.0 | 849 | 0.006 |
|  |  | 1 | 970 |  | 55.3 | 2,352 | 0.044 |
| Czech Rep. & SE Germany | 53,670 | 2 | 4959 | 11 | **10.8** | **460** | **0.009** |
|  |  | 3 | 14826 |  | 3.6 | 154 | 0.003 |
|  |  | 1 | 3160 |  | 7.3 | 310 | 0.013 |
| Switzerland & SW Germany | 23,050 | 2 | 4336 | 14 | **5.3** | **226** | **0.010** |
|  |  | 3 | 9988 |  | 2.3 | 98 | 0.004 |
|  |  | 1 |  |  | 10.3 | 437 | 0.013 |
| Italy | 32,500 | 2 |  |  | **7.5** | **319** | **0.010** |
|  |  | 3 |  |  | 3.3 | 138 | 0.004 |
|  |  | 1 |  |  | 6.2 | 262 | 0.013 |
| SE France | 19,480 | 2 |  |  | **4.5** | **191** | **0.010** |
|  |  | 3 |  |  | *2.0* | *83* | *0.004* |
|  |  | 1 | 2420 |  | *10.2* | *434* | *0.018* |
| SW France | 24,740 | 2 | 4337 | 10 | **5.7** | **242** | **0.010** |
|  |  | 3 | 13094 |  | *1.9* | *80* | *0.003* |
|  |  | 1 |  |  | *31.3* | *1329* | *0.018* |
| Spain & French Pyrenees | 75,710 | 2 |  |  | **17.5** | **742** | **0.010** |
|  |  | 3 |  |  | *5.8* | *246* | *0.003* |
|  |  | 1 |  |  | *3.6* | *154* | *0.018* |
| Portugal | 8,780 | 2 |  |  | **2.0** | **86** | **0.010** |
|  |  | 3 |  |  | *0.7* | *28* | *0.003* |
|  |  | 1 |  |  | **207** | **8794** | **0.018** |
| **Sum Core Area** | **480,560** | 2 |  | **39** | **100** | **4254** | **0.009** |
|  |  | 3 |  |  | **52** | **2209** | **0.005** |
|  |  | 1 |  |  |  |  | **0.003** |
| **Total Area of Calculation** | **2,600,000** | 2 |  |  |  |  | **0.002** |
|  |  | 3 |  |  |  |  | **0.001** |

For the <u>estimates from dataset B</u> (see **S3 and S4 Tables**), the size of all Core Areas of GI-1d-a (**Fig 3**: shaded areas) is reduced to 489,000 km^2^ (85% of the sites; 842 out of 988), and those of GS-1 to 392,000km^2^ (**Fig 4**: shaded areas; 74% of the sites; 415 out of 560). The mean estimate for the total population for GI-1d-a sums up to only roughly 5700 people (min. 3100, max. 9900) and to 3500 people (min. 1900, max. 6200) for GS-1. The mean density of GI-1d-a, 1.2 p/100 km² (min. 0.6, max. 2.0), in contrast, is only slightly lower than for dataset A, and remains at 0.9 p/100 km² for GS-1 (min. 0.5, max. 1.6). The largest regional population for dataset B estimates during GI-1d-a is found in northern Spain and the French Pyrenees with 1100 people, followed by Benelux and NW Germany with 820 people. With 770 people, NE Germany and Poland display estimates similar to those of the Czech Republic and SE Germany.

The largest differences between estimates from datasets A and B are observed during GI-1d-a especially for the Czech Republic and SE Germany, but also for the area of Poland and NE Germany and for Switzerland and SW Germany (**Fig 5**, **for details, see S5 Fig**). For GS-1, differences are observable for the Czech Republic and SE Germany, where dataset A estimates around 460 people and dataset B around 80 people (**S6 Fig**). The density for this area, however, remains at 2.2 p/100 km² for GI-1d-a and 0.9 p/100 km² for GS-1, fully in line with the dataset A estimates. For the total area of calculation the estimated population density drops in dataset B from 0.002 in GI-1d-a to 0.001 p/km^2^ in GS-1.

**Fig 5.**
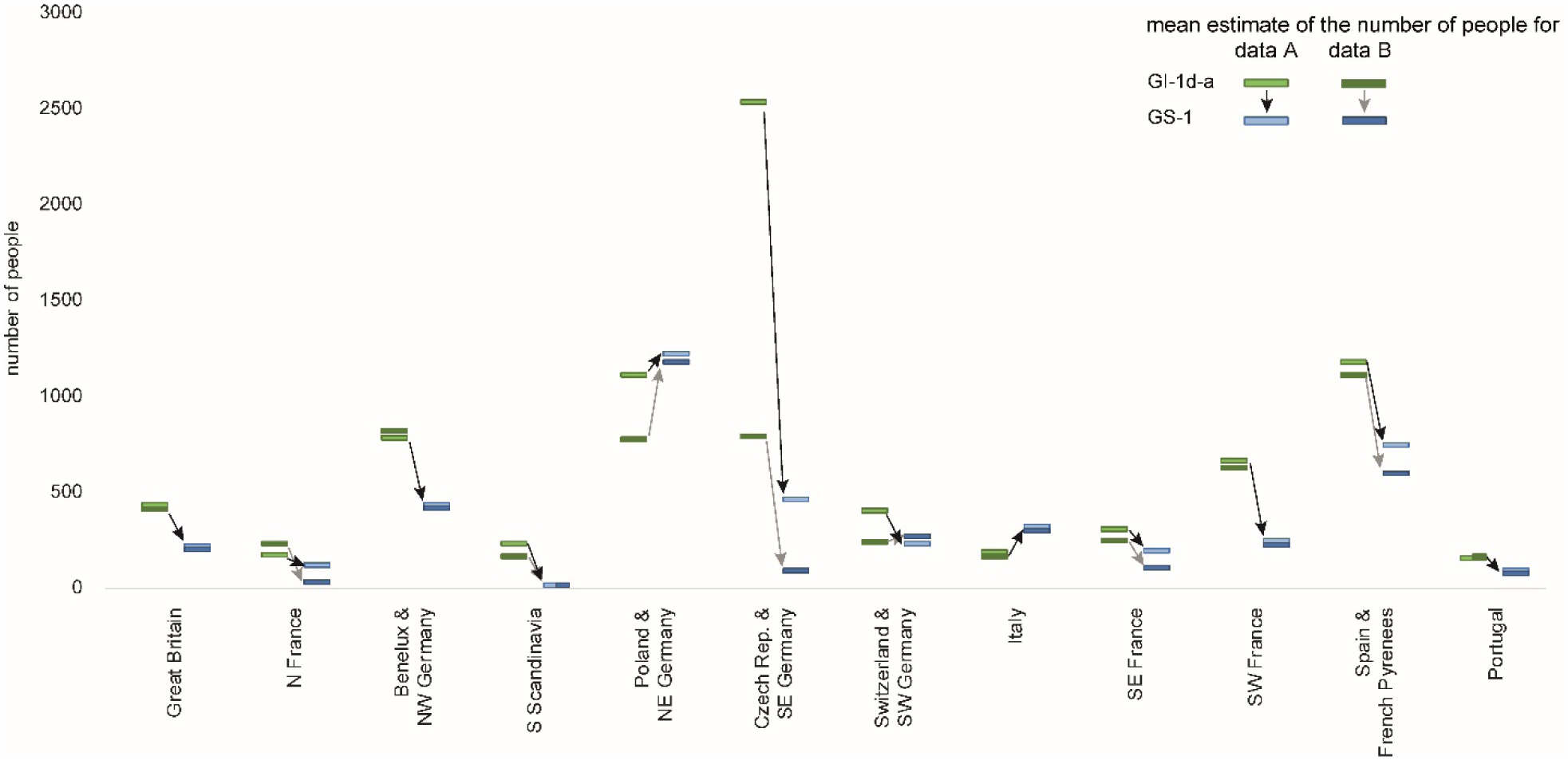
Comparison of demographic estimates from GI-1d-a and GS-1. Depicted are the mean estimates from dataset A (bright fill) and dataset B (dark fill).

A consistent signal in our palaeodemographic reconstructions is a strong decline in the meta-population from GI-1d-a to GS-1 (**Fig 5**). In Portugal, SE France, and S Scandinavia, likewise Czech Republic and SE Germany, the regional population may have dropped below a hundred people. Notable exceptions from this trend are Poland, NE Germany, and Italy (especially the northeastern area, compare **Figs 4** and **5**). Moreover, in all estimates, the former region hosts the largest regional population of around 1200 during GS-1, thus being almost twice as large as the second highest regional population (data A = 740; data B = 590) located in (northern) Spain and the French Pyrenees. In Switzerland and SW Germany, the estimates from dataset B result in a relatively stable population (data A = 260, data B = 230).

## Discussion

Large-scale distribution patterns of Palaeolithic sites are influenced by several confounding factors that lie outside the decision-making processes of past hunter-gatherers. Many other parameters can influence the likelihood of site discovery, such as the geomorphological development of a landscape as shown, for instance, in the Great Adriatic Po-Plain [123] and other coastal areas [124]. Modern land use (e.g., forestry, vineyards, ploughed fields, quarries) or specific taphonomic settings (presence of caves and rock shelters or wetlands; loess or coversand deposition) can also play an important and sometimes even dominant role [125]. Other factors are the research histories of the different areas, in some cases linked to the proximity of universities and other institutes (research-driven versus development-led archaeological excavations) with strong prehistoric research traditions, the accessibility of collected data (such as regional or national databases, or grey literature) and their revision by experts (cf. [67,109,126,127]). Finally, differences in artefact size or the recognisability of iconic artefacts, such as the tanged points of the Ahrensburgian and Swiderian, can skew the observed patterning, either by making them harder to find (cf. [73,128]), or by skewing site numbers towards those characterised by particularly large artefacts (cf.[129]). These biases need to be discussed and necessitate caution but should allow research to address large-scale questions such as population dynamics in general [15].

The distribution of Final Palaeolithic sites in the study area (**Fig 2**) is notably uneven but comparable between GI-1d-a and GS-1 (**Figs 3 and 4**). Generally – and in sharp contrast to all periods of the preceding Upper Palaeolithic – the main area of the human population is not located in the Franco-Cantabrian region but in regions north of the Alps. Given that modern land use will not affect Upper and Final Palaeolithic sites differently and given further that the Late Glacial warming made higher-latitude areas newly accessible for Final Palaeolithic hunter-gatherers, it seems highly unlikely that this large-scale spatial pattern is exclusively the result of research biases [125]. At smaller scales, however, the effects of research biases might be more pronounced. For our dataset, this is particularly relevant for a conspicuous void that stretches in the south-west to north-east direction from southwestern France up to the Upper Rhine Valley, separating the investigated area into a south-western and north-eastern part. Final Palaeolithic sites are scarce in this void, but it is unclear whether this reflects prehistoric reality or results from multiple biases listed above. The latter view is supported by the absence of strong research traditions and academic structures in this region compares with, for instance, the southwest or the northeast of France and a relative scarcity of contexts favouring preservation and recognition of Upper and Final Palaeolithic sites. Difficulties in locating raw material sourcing sites and a general scarcity of typological markers (e.g., at Le Closeau) might additionally hamper the attribution of sites to the Final Palaeolithic in some areas. Also, modern population densities and industrial activity are either low or very high in some of the western areas of Germany – some sites are surely waiting to be discovered while others have long since been destroyed. The comparably long and strong archaeological research tradition here, however, speaks in favour of the existence of a void, at least in Germany (i.e. Hesse and western Thuringia). A comparison of the extent of the void with Core Areas calculated for the immediately preceding period of the Final Magdalenian (**Fig 6**, [122]), shows that the void widens considerably in the Final Palaeolithic towards the southwest and northeast. Core Areas are modelled in the void, at least for the French area, and also during earlier periods of the Upper Palaeolithic record, the early Gravettian and Solutrean technocomplexes [50,130]. It is difficult to see why potential biases would particularly affect Final Palaeolithic sites, leaving those of the Upper Palaeolithic in place. Also, differences in material culture on either side of the void (e.g. occurrences of painted cobbles), a lack of evidence for objects that were transported across it, and hints from palaeogenetic data warrant a closer examination of this observation.

**Fig 6.**
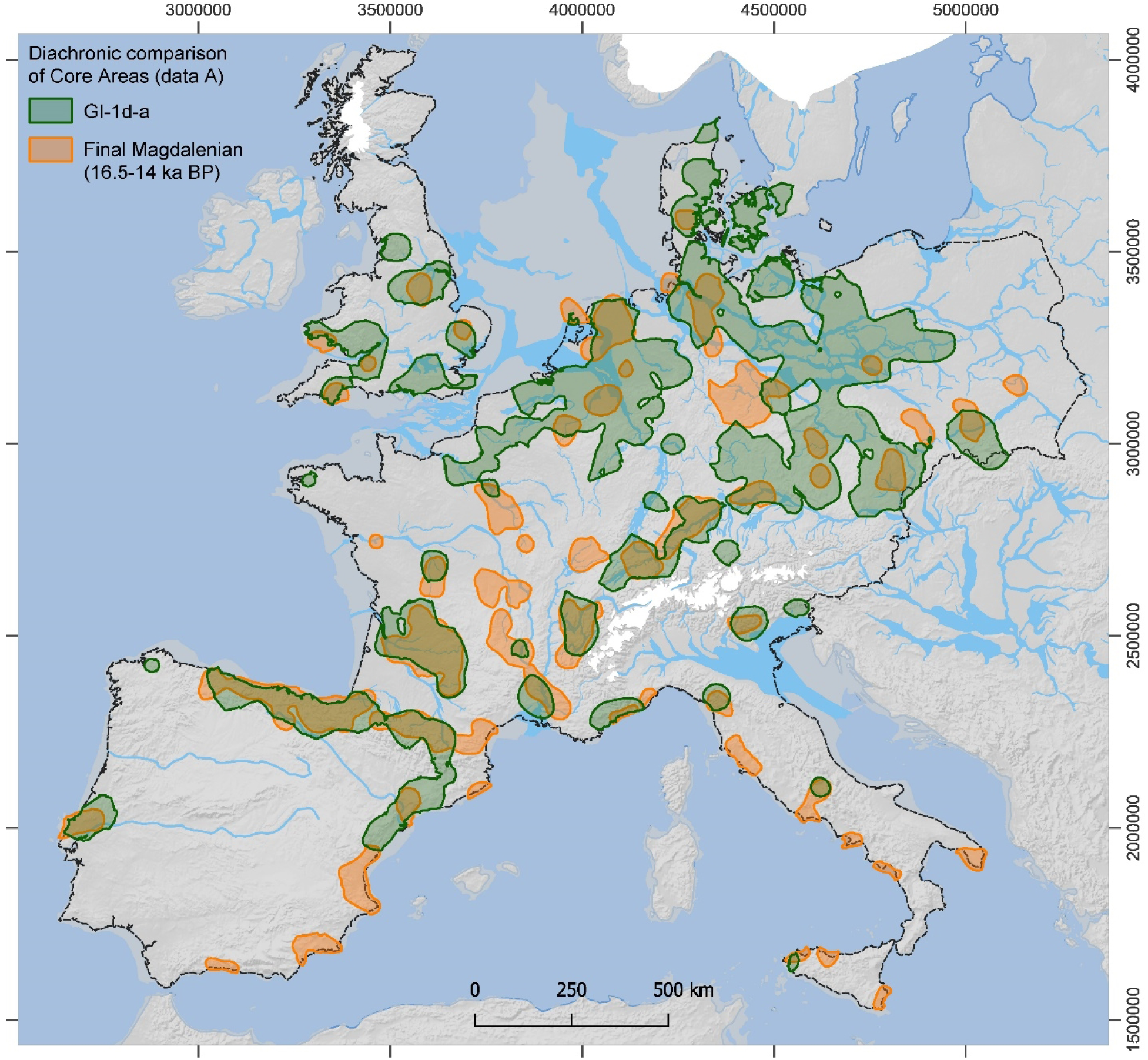
Comparison of Core Areas from the Final Magdalenian (ca. 16-14 ka BP, orange areas, after [122]) to Core Areas of the GI-1d-a (green areas). For details on background map, see Fig 3.

In order to assess biases by typological visibility, we had a look at the number of radiocarbon-dated sites in both phases based on the entries in the Radiocarbon Palaeolithic Europe Database v.27 ([131]; we are aware of a later version but used v.27 when conducting this study). The analysis shows similar trends to our demographic estimates. For the entire map section and in absolute numbers, there is a decline of more than 50 % in radiocarbon dated sites from GI-1d-a to GS-1 (**S5 Table**). Regionally differentiated and in relative terms (**S7 Fig**), we see a decline in all areas, but it is specifically pronounced in Central Europe. A fundamental difference, however, is our increasing estimates for NE Italy, Poland, and NE Germany (**Fig 5**). It is thus possible that our estimates for these regions are positively skewed due to greater typological recognisability of assemblages produced during GS-1. The estimated demographic increase, however, can also be the result of behavioural responses to contemporaneous climate change, such as population movements towards the east.

A comparison to other palaeodemographic studies might also reveal potential biases. Yet, in contrast to periods of the Upper Palaeolithic, such a comparison is difficult for the Final Palaeolithic estimates because of divergent chronological frameworks. For instance, the youngest period in the study by Bocquet-Appel et al. [132], is termed ‘Late Glacial’ and comprises the interval roughly between 19.1 and 13.3 ka BP. Tallavaara et al. [20] provide estimates in 1 ka steps, but the last two steps are centred at 14 and 13 ka BP, respectively, therefore essentially missing the Younger Dryas. While the former period is too large for a meaningful comparison, the latter provides two estimates at the fringes for our first phase and none for the second phase of GS-1. The chronologically matching study by French and Collins for southwestern France [19] also sees a decline in demographic signatures from GI-1d-a to GS-1. A revision of the Final Magdalenian assemblages in this area resulted in a reattribution to the Laborian, chronologically placing the sites to the GS-1 [133]. These findings and very recent research developments on the Laborian could change the demographic picture here. Rather more synoptically on a European scale, Ordonez and Riede [134] reconstruct forager population densities with reference to ethnographic data and climatic limiting factors as derived from high-resolution climate data. Their study similarly documents a population downturn during GS-1.

### A large-scale shift to the east as a potential reaction to the GS-1 cooling

At a large spatial scale, a ∼25% decrease in the size of the Core Areas from GI-1d-a to GS-1 is observable in most parts of the investigated area and across both datasets A (**Fig 7**) and B (**S8 Fig**). At the same time, Core Areas in western Britain, Brittany, and Denmark disappeared, whereas new Core Areas formed in eastern Poland, particularly in Masuria. Core Areas in Iberia and southwestern France shrank but remained rather stable in their geographic location. Notably, in the rest of the investigated area, Core Areas showed a pronounced geographic shift towards the east during GS-1. Given that it seems unlikely that this large-scale shift is exclusively caused by biases (see discussion above), the question arises as to whether it can be seen as a reaction to changing climatic conditions. Here, a comparison to the developments during the Gravettian is instructive. Demographic studies, which compared the internal shift from an early phase (33-29 ka BP) to a late phase (29-25 ka BP) of the Gravettian is the only other instance since the arrival of anatomically modern humans in Europe where we see a pronounced demographic decline coinciding with climatic cooling [50]. During the late phase of the Gravettian, the estimates indicate a reduction across all regions, suggesting that no or only little population movement occurred over larger distances. This may indicate that a behavioural response to environmental change in terms of mobility did not happen or that population movement was not a sufficiently effective risk mitigation strategy, given that the climatic changes affected all of Europe [135]. Instead, it seems that the regional populations north of 50° latitude collapsed entirely, leaving western Central Europe largely depopulated until the Late Magdalenian [50,136]. This is in accord with palaeogenetic findings, indicating a continuity in western and southwestern Europe (Fournol cluster) and a disappearance of traits associated with the Věstonice cluster in Central and Southern Europe [23]. Interestingly – and in stark contrast to the Gravettian – the estimates for the Final Palaeolithic show a regionally more differentiated demographic pattern. In the following, we discuss the environmental change of the Younger Dryas in more detail, focusing on regional differences that might have contributed to the observed pattern.

**Fig 7.**
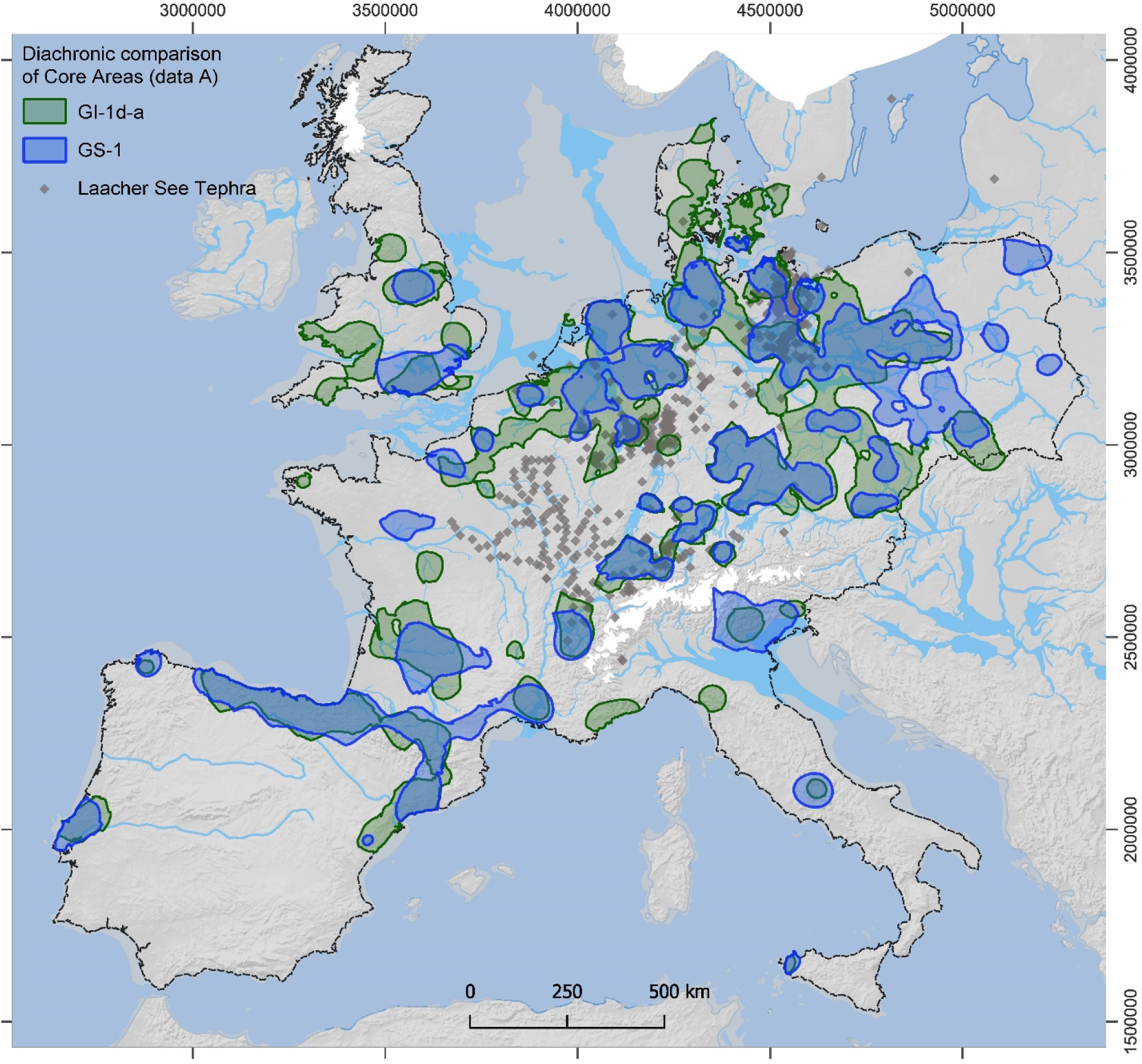
Comparison of Final Palaeolithic Core Areas (based on data A) from GI-1d-a to GS-1. For a comparison of data B Core Areas, see **S8 Fig**. Grey diamonds (after [137]) indicate documented ash (=tephra, microtephra or heavy minerals) from the Laacher See eruption, dated to 13 ky BP [138].

The Younger Dryas (GS-1) is a phenomenon driven by processes in the North Atlantic and with partly varying effects on the ecosystems in different parts of Europe, often with a west-east-trending gradient (e.g., [34–37,41,109,139,140]). In northern Germany, western Belgium [141] and Poland [142], for instance, the climate of GS-1 drove the return of open landscapes, while the vegetation in Switzerland and eastern France seems to have been affected only weakly [143–145]. In a general comparison to the GI-1d-a, there is a sharp decline in winter temperatures with much colder temperatures between September and May, with strongly increased snow depth in north-western Europe during May, particularly in western Britain, Brittany, and Central France and a lesser extent in northern Central Europe [41].

As a consequence of this marked cooling, the thawing of the permafrost, ice-covered lakes, and other bodies of water was delayed. The period with stable warm summer temperatures was considerably shorter, restricted to about three months (June to August) in eastern Europe and to only two months (July to August) in north-western Europe (e.g., [40,146,147]). This led to a considerable decline in the length of the growing season, again with a pronounced west-east trend. While the regions along the Atlantic coast of France and Britain experienced a shortening of more than a month, Eastern Britain, Central France, Iberia, and Italy saw a reduction of 10-20 days, while northern Central Europe and the Balkans saw a decrease of between 2 and 10 days [41]. Shifts in the phenological patterns of the growing season can have considerable influence on the direction of migrations of herbivores [148] and might, over the course of centuries also influence the large-scale distribution of hunter-gatherer populations [22,149]. Interestingly, isotope analyses on reindeer teeth from the Hamburgian and Ahrensburgian layer at Stellmoor (northern Germany) suggest that reindeer herds generally migrated in an east-west direction on the North European Plain during the Younger Dryas [150], although these patterns could have been more complex in detail [151], cf. [38]. Another highly relevant phenomenon is a general shift towards dryer conditions during the Younger Dryas, which probably led to the lowering of the groundwater table and, thus, reduced water availability in some areas [152,153]. This phenomenon is particularly well documented in the Scheldt basin in north-western Belgium. Here, a lowering of the groundwater table probably led to a disappearance of numerous shallow lakes and ponds in the interior of the Lower Scheldt basin, while the neighbouring Meuse basin seems to have been less affected [140]. In addition, the lowlands of the North European Plain were affected by intensified winds forming cover sand dunes [154,155]. Lower groundwater tables, sand cover in which water seeps away easily and longer periods with ice-cover of the remaining water bodies might have posed challenges to animals and humans alike. Yet, the cover increased the in situ preservation of sites but also reduced their archaeological visibility [156].

In sum, the Younger Dryas has precipitated a swath of largely unfavourable environmental changes for humans. These changes were more pronounced closer to the Atlantic seaboard and less so towards the east. This is in line with our observation of an eastward shift of the Core Areas in north-western and Central Europe. While populations declined in most areas, there is a clear increase in NE Germany and Poland as well as stable estimates for Switzerland and SW Germany in the scenario based on dataset A. At the same time, the estimates for SE Germany and the Czech Republic show the sharpest decline of all regions in both scenarios, dataset A and B. In S Scandinavia, i.e. Denmark, the population dropped almost to zero, with only a few camps appearing in the East. Considering the overall decline of regional populations, independent population growth in NE Italy, NE Germany, and Poland seems unlikely. Therefore, these regional population dynamics may be seen as first evidence of large-scale population movement as a response to climatic cooling. If the population shift represents a prehistoric reality, then the source of the movement into NE Germany and Poland would have mainly been SE Germany and the Czech Republic, with some people moving towards Switzerland but the majority moving towards Poland. The sink areas of Poland, NE and NW Germany also could have had an influx from Denmark. NE Italy might have received an influx from SE France or western Italy, although there is a lack of archaeological evidence for contact in this direction [157]. Possible source areas are also regions at the margins or outside the investigated area, such as central-southern Italy, the western Balkans, or now submerged coastal areas.

Finally, environmental events beyond climate trends may also have played a role in shaping human responses at this time. The spatial congruence between the void area across Central Europe and the most massive ash fallout of the cataclysmic Laacher See eruption that occurred at 13,006±9 cal BP [138] in the Eifel volcanic zone is notable, especially to the south-east and north-east of the eruptive centre (**Fig 7**). The eruption occurred at the tail end of the cool GI-1b which likely stressed contemporaneous forager populations. In Belgium and Central Germany, several sites evidence stratigraphic relations where *Federmesser-Gruppen* occupation is capped by Laacher See fallout, terminating these occupations [158,159]. The short-and medium-term environmental impacts of the eruption were varied and, depending on local fallout thickness, likely considerable [160–164]. These may have depressed population in the affected regions already prior to the onset of GS-1 and may have resulted in sustained legacies of landscape avoidance that lasted into GS-1.

### Demographic dynamics, minimal viable population, and social networks

Depending on whether one favours the mean estimate of the total population during GI-1d-a from dataset A (8100, also including generically assigned sites) or the one from dataset B (5700, using only chronologically subdivided sites) as more reliable, there is either an increase (dataset A) or decrease (dataset B) after the Final Magdalenian, which produced a mean estimate of 7300 (min. 4500, max. 10200) people [122]. However, some northern regions were frequently occupied for the first time after the Last Glacial Maximum (**Fig 6**), suggesting a stable metapopulation if not an increase. Less ambiguous is the subsequent decline in GS-1 to 4300 and 3500 people for the estimates based on datasets A and B, respectively. In both scenarios, the population is reduced by roughly half. This marked decline raises questions about the viability of the remaining population and the consequences for regional as well as long-distance social networks.

Defining what constitutes minimum viable populations in humans is difficult. A lower threshold of ∼1500 individuals, comprising around 770 individuals in the reproductive age cohort, has been suggested [165–167]. While the effects of drift-related inbreeding can be avoided at even lower numbers of 100 to 500 people [166,168], such minuscule populations become highly sensitive to negative stochastic effects, especially in high-risk environments [169]. Despite being relatively low, our estimates are in the range of viable populations, including, notably, the estimates for GS-1, which are still high compared to those of most other periods of the Upper Palaeolithic [130,170]. This difference is particularly striking regarding the Gravettian, where mean estimated population sizes dropped from 2800 people during the early phase to only 1000 during the late phase [50].

The estimated population decline likely also affected the size, structure, and connectivity of social networks. On the one hand, a reduction in the number of people who can maintain social contacts is likely to lead to distortions in the networks. Here, networks at regional scales are usually maintained by more people than networks at larger scales, which are thus more prone to disturbances. On the other hand, actively increasing the extent and connectivity of networks can be a way to mitigate environmental risk and population decline [171] – as long as the increasing physical distances can be bridged (cf. [172–174]). Social networks exist at different spatial scales, and besides palaeogenetic studies [23], they usually become archaeologically visible through different classes of objects circulating within them that have characteristic transport distances and patterns [175]. For networks at a regional scale, lithic raw materials are probably the most important source of information [176]. The materials usually transported in economically significant numbers and over distances well below 300 km are here considered likely to be acquired by embedded procurement [177,178] and thus by members of the same regional group that also discarded them (but see [179]). In the available raw material data, there is indeed an increase in the size of the catchments during the Final Palaeolithic, very pronounced in all areas of the study region except for the NW European area (see RMCA median values in **Tables 1 and 2**; **S1 and S2 Tables**).

For large-scale networks (connecting two or more regional networks) to become visible, transport patterns of objects with a presumably different acquisition mode are better suited. Ideally, these objects are not procured directly by the group that discards them but acquired via exchange between groups. Here, sub-recent and fossil mollusc shells seem to be highly informative. In contrast to lithic raw materials, they are regularly transported over more than 600 km and during the Late Upper Palaeolithic, they trace a network that spans between the Atlantic and Mediterranean coast, the Paris Basin up the Rhine-Meuse Area and across to the Swabian Jura [180–183]. They thus also span across the area of the central void in the Final Palaeolithic dataset (**Fig 7**). For the Final Palaeolithic itself, however, information on transported mollusc shells is extremely rare. There are singular instances of long-distance transports of lithic raw materials (e.g. obsidian found in Poland), yet there is no available evidence that systematically reflects social networks at that scale. The absence of evidence for regular long-distance transports of objects in contrast to the Late Upper Palaeolithic may signal reduced contact at that scale. By this token, the extensive void in the centre of the investigated area might also be an expression of reduced social connectivity between hunter-gatherers in southwestern France and Iberia on the one hand and in Northern and Central Europe on the other. The latter area can possibly be further divided by mid-scale networks through the distribution of raw materials such as red Helgoland flint [52] or the use of amber [184,185] or by differentiated symbolic use [97,185]. If there had been decreasing contact between the south-western and north-eastern parts of the investigated area during GI-1d-a, the marked population decrease in GS-1 probably would have additionally distorted the networks.

Recent palaeogenetic evidence suggests a differential development in both regions during the Final Palaeolithic. There appears to be a probably south-north directed movement of people during GI-1d-a, passing west of the Alps from Italy into Central Europe and the British Isles, whereas the population in Iberia shows more pronounced genetic continuity with several admixture events [23,66]. Whether this suggested south-north dispersal is consilient with the proposed northward dispersal of backed points from south-eastern France along the Rhône and Saône axis and alongside the disappearance of reindeer, as often emphasised by Thévenin [45,186,187], is beyond the scope of this paper. However, these proposals have mainly been based on typological analyses carried out on assemblages whose archaeostratigraphic reliability is difficult to prove. Furthermore, given the available data, the Azilian traditions appear after the Magdalenian, and some can be considered as a final episode of these traditions [3,90,188]. Other, more recent traditions of the Azilian (GI-1) are part of their extension and of a common evolution of technical traditions on a European scale. The situation is perhaps more complex at the very end of the Palaeolithic (GS-1) with the existence of a certain duality of technical traditions in France. The presence of assemblages with characteristics close to those known from the Laborian and Epigravettian periods has been proposed [83,93,96,189]. However, the chronological framework of these phenomena remains to be defined and the techno-economic data needs to be improved. At the moment, identifying this large empty area with very little to no evidence being assignable to the Final Palaeolithic calls for future research.

## Conclusion

During the GI-1d-a, the higher-latitude areas of Europe – parts of which were previously unoccupied or only ephemerally visited by anatomically modern humans – became more regularly and frequently visited by contemporaneous hunter-gatherer groups. At this time, the western and central European lowlands became a hotspot of population dynamics for the first time. The climatic cooling during GS-1 (Younger Dryas) led to a marked population decline but also different regional responses. Populations in the Iberian Peninsula and southwestern France shrank while staying in the same area, a response comparable to the one observed during the Gravettian. Populations in Northern and Central Europe, in contrast, reacted with a pronounced shift towards the east. While populations declined in the western part, they increased in the eastern part. We argue this was not because of population growth but because of population movement from west to east. The same phenomenon can be observed for the area of south-eastern France up to north-eastern Italy, where the former region also showed a population decline, while the latter saw a population increase, again possibly due to population influx. The shift of populations towards the east was most probably a reaction to deteriorating environmental conditions brought about by the Younger Dryas, including longer winters, higher snow cover and a strongly reduced growing season. These changes affected the western parts of Europe particularly hard, especially the Atlantic coast, and became more and more buffered towards the east.

The question thus arises as to why the two sub-populations in the southwestern part and the rest of the investigated area reacted so differently. Answers to this question are not readily evident. Maybe the two sub-populations had already become largely decoupled during the Allerød, as might be indicated by the central void area. Maybe hunter-gatherers in the topographically rather uniform lowlands had a different attitude towards and aptitude for long-distance movements or a more permeable territorial structure. The possibility that some behavioural or technological innovation – using poison for efficiently killing prey, snowshoes, skis, travois or the like – transformed their ability to move longer distances at lower costs should not be discounted. Future studies are needed to illuminate these aspects. To further refine our palaeodemographic estimates, a higher chronological resolution of the dataset will likely reveal currently invisible patterns, which in combination with high-resolution climate models and palaeoenvironmental studies will allow for a more differentiated view of the biocultural processes that occurred at the end of the Pleistocene. In-depth studies of material culture change and a better understanding of transport patterns of raw materials and objects of far-distance transports could provide data on the size and connectivity of social networks and therefore offer additional details to the coarse picture and potentially identify non-environmental explanations for the observed patterns of population dynamics.

## Acknowledgements

The authors would like to thank Andrea Darida, Martin Müller and Nina Avci for their help in compiling the database and Dennis Batz for editing the references. All authors revised and added new data on sites and assemblages for regions of their expertise. Felix Riede, Katja Winkler, Florian Sauer and Birgit Gehlen provided regional databases at an early stage of the project, that were incorporated after adjustment to the specific requirements. All geospatial calculations were conducted using MapInfo 8.5, maps were created using QGis 3.34. The German Research Foundation funded the research of Isabell Schmidt – Project-ID 57444011 – SFB 806 ‘Our Way to Europe’. Andreas Maier’s contribution is made as part of the project ‘HESCOR’, funded from the programme ‘Profilbildung 2022’, an initiative of the Ministry of Culture and Science of the State of Northrhine Westphalia (HESCOR PB22-081). Research contributed to this paper by Berit Valentin Eriksen, Mara-Julia Weber and Sonja B. Grimm was funded by the DFG, German Research Foundation, project number 290391021-SFB 1266, ‘Scales of Transformation’. Research of Werner Schön in southwest Bavaria (Allgäu) was funded by the DFG, German Research Foundation, project numbers 386654307 and 57444011. Felix Riede’s contribution is made as part of CLIOARCH, an ERC Consolidator Grant project, and has received funding from the European Research Council (ERC) under the European Union’s Horizon 2020 research and innovation programme (grant agreement No. 817564). Annabell Zander’s contribution is made as part of her British Academy Postdoctoral Fellowship at the University of York. MN would like to thank Dion Stoop (Noordelijk Archeologisch Depot, Nuis), Huub Scholte Lubberink (RAAP Oost, Zutphen) and Haije Veenstra (RAAP Noord, Drachten). The authors are solely responsible for the content of this publication.

## Supporting information

### Information on Maps

#### Tables

S1 Table. Sites with data on Raw Material Catchment Areas (RMCA) assigned to GI-1d-a. S2 Table. Sites with data on Raw Material Catchment Areas (RMCA) assigned to GS-1.

S3 Table. Regionally distinguished demographic estimates for GI-1d-a based on dataset B (i.e., only directly dated or specifically attributable to GI-1d-a).

S4 Table. Regionally distinguished demographic estimates for GS-1 based on dataset B (i.e., only directly dated or specifically attributable to GS-1).

S5 Table. Number of radiocarbon-dated sites divided by GI-1d-a and GS-1.

#### Figures

S1 Fig. Maps (left) and diagrams (right) from the definition of the Core Areas based on the sites dated to Greenland Interstadial (GI) 1d-a.

S2 Fig. Maps (left) and diagrams (right) from the definition of the Core Areas based on the sites dated to Greenland Stadial (GS) 1.

S3 Fig. Map of all Raw Material Catchment Areas (RMCA) from sites dated to GI-1d-a that were collected from the literature (n = 108).

S4 Fig. Map of all Raw Material Catchment Areas (RMCA) from sites dated to GS-1 that were collected from the literature (n = 97).

S5 Fig. GI-1d-a: Comparison of estimates of people for dataset A (left bar) and B (right bar) per region.

S6 Fig. GS-1: Comparison of estimates of people for dataset A (left bar) and B (right bar) per region.

S7 Fig. Diachronic tendencies in the frequency of regionally differentiated radiocarbon dates per region.

S8 Fig. Comparison of Final Palaeolithic Core Areas (data B) from GI-1d-a (green areas) to GS-1 (blue areas).

**Supporting information References**

