## Supplemental file for "Demographic responses to climatic changes during the Final Palaeolithic in Europe"

### Supporting information for: Demographic responses to climatic changes during the Final Palaeolithic in Europe

### Information on Maps

All maps were created using QGIS, version 3.34.7 “Prizren” in projection WGS 84, EPSG 3035.

The calculation of the palaeodemographic estimates, including the modelling of the optimally describing isoline / Core Areas (see [1] and https://github.com/C-C-A-A/CologneProtocol-MapInfo) and the convex hulls of the raw material catchment areas (RMCA), was conducted using MapInfo Professional 8.5, projection Lambert azimuthal Flächentreu - Baltikum (Bereich 90).

The GIS-sources used to create the basemap were taken from https://www.naturalearthdata.com/:

Relief shading and hypsography:

1:10m Gray Earth: Gray Earth with Shaded Relief and Hypsography

Coastline / Ocean:

1:50m Physical Vectors, Ocean

Rivers and Lakes

1:50m Rivers, Lake Centerlines

Additionally, we included for areas > 45° N:

Open access data from EPHA – European prehistoric and historic atlas (https://zbsa.eu/european-prehistoric-and-historic-atlas/):

- Drainage Systems
- Allerød
- Dryas III

The GIS-created maps were reworked and a legend was added in Adobe Illustrator CS6.

### Tables

#### S1 Table. Sites with data on Raw Material Catchment Areas (RMCA) assigned to GI-1d-a.

| Region | Site | RMCA in km$^2$ | Reference |
|---|---|---|---|
| NW | | | |
| | Bad Breisig | 549 | [2]; pers. comm. B. Gehlen |
| | Eindegoorheide 1 | 630 | [3] |
| | Geldrop 3-1 | 797 | [4] |
| | Urbar (20) | 853 | [5] |
| | Tongeren-Plinius | 2.670 | [6] |
| | Wesseling-Eichholz | 3.665 | [7,8]; pers. comm. B. Gehlen |
| | Horn-Haelen | 3.909 | [9] |
| | Gönnersdorf | 4.959 | [10] |
| | Heythuysen-De Fransman | 5.117 | [9] |
| | Kettig | 5.220 | [11] |
| | Andernach-Martinsberg | 5.818 | [12]; pers. comm. B. Gehlen |
| | Rüsselsheim 122 (A, B) | 5.896 | [13] |
| | Niederbieber II | 7.284 | [12]; pers. comm. B. Gehlen |
| | Niederbieber IV | 7.284 | [12]; pers. comm. B. Gehlen |
| | Ruien "Rosalinde" | 7.562 | [14] |
| | Kartstein | 9.665 | [15]; pers. comm. B. Gehlen |
| *NW* | *Q1* | *2.670* | |
| | *Q2* | *5.117* | |
| | *Q3* | *5.896* | |
| NE | | | |
| | Calowanie (Calowanie) | 605 | [16] |
| | Jeskyně tří volů (Jeskyne tri volu) | 747 | [17] |
| | Tarnowa 1 | 975 | [16] |
| | Dzierżysław (Dzierzyslaw) | 1.302 | [18] |
| | Kvíc | 1.767 | [19] |
| | Kraków-Bieżanów (Krakow-Biezanow) | 2.129 | [20] |
| | Trebomyslice 1 | 2.318 | [21] |
| | Hostim I - Krápníková jeskyne | 2.729 | [17] |
| | Uckersdorf | 24.016 | [22] |
| | Wöllershof-Ketzerrang | 24.923 | [22] |
| *NE* | *Q1* | *1.057* | |
| | *Q2* | *1.948* | |
| | *Q3* | *2.626* | |
| SE | | | |
| | Abri de Douattes; Douattes Est | 689 | [23] |
| | Sattenbeuren-Kieswerk | 876 | [24] |
| | Gunzwil-Beromünster | 1.523 | [23] |

| Region | Site | RMCA in km² | Reference |
|---|---|---|---|
| [SE] | Wauwil-Sandmatt 25 | 1.882 | [25] |
| | Birseck-Ermitage | 2.352 | [23] |
| | Abri Unter den Seewänden | 2.857 | [26]; pers. comm. B. Gehlen |
| | "Gemeindebeunden" Bad Buchau-Kappel | 3.056 | [27] |
| | Neumühle, Abri / Neumuhle | 3.110 | [23] |
| | Geispel | 3.474 | [23] |
| | Monruz (Neuchâtel) | 3.550 | [23]; pers. comm. J. Affolter / B. Gehlen |
| | Fru, La (423) | 3.864 | [28] |
| | Champreveyres / Hauterive-Champréveyres | 5.023 | [23] |
| | Cure, Abri de la | 7.467 | [23]; pers. comm. J. Affolter / B. Gehlen |
| | Lüscherzmoos | 8.297 | [23]; pers. comm. J. Affolter / B. Gehlen |
| | Cham-Grindel III | 9.026 | [23] |
| | Langrüti | 13.833 | [23] |
| | Fürsteiner | 15.239 | [23] |
| | Schlössel 1 | 20.434 | pers. comm. J. Affolter / B. Gehlen |
| | Hopferau-Pertlesbichl | 28.503 | [29]; pers. comm. B. Gehlen; |
| *SE* | *Q1* | *2.604* | |
| | *Q2* | *3.550* | |
| | *Q3* | *8.662* | |
| SW | | | |
| | Chaloignes, Les | 571 | [30] |
| | Forcas I | 755 | [31] |
| | Balzola, Cueva de | 2.401 | [32] |
| | Troubat, La Grotte-Abri de/ Moulin | 2.522 | [33] |
| | Rhodes II, L'Abri | 12.788 | [34] |
| | Béraud, Grotte | 28.672 | [35] |
| | Mas d'Azil | 40.446 | [36] |
| *SW* | *Q1* | *1.578* | |
| | *Q2* | *2.522* | |
| | *Q3* | *20.730* | |

The table comprises only data that were considered for calculation in this study (n = 53). Below each region, the corresponding quartiles (Q1, Q2, Q3) used to calculate the palaeodemographic estimates are shown (see **table 1**).

#### S2 Table. Sites with data on Raw Material Catchment Areas (RMCA) assigned to GS-1.

| Region | Site | RMCA in $km^2$ | Reference |
|---|---|---|---|
| NW | | | |
| | Geldrop 3-1 | 797 | [4] |
| | Altenrath-Ziegenberg | 3.646 | [12]; pers. comm. B. Gehlen |
| | Reingsen I | 6.728 | [37–39]; pers. comm. B. Gehlen |
| | Kartstein | 9.665 | [15]; pers. comm. B. Gehlen |
| *NW* | *Q1* | *2934* | |
| | *Q2* | *5187* | |
| | *Q3* | *7462* | |
| NE | | | |
| | Calowanie (Calowanie) | 605 | [16] |
| | Chełmno 4 (Chelmno) | 963 | [40] |
| | Rzuchów 24 | 963 | [40] |
| | Janów 21 | 976 | [40] |
| | Dzierżysław (Dzierzyslaw) | 1.302 | [18] |
| | Gönnersdorf | 4.959 | [10] |
| | Cichmiana 2 | 7.924 | [40] |
| | Fürth-Atzenhof-NO | 11.408 | [22] |
| | Blanice 6 | 18.244 | [19] |
| | Uckersdorf | 24.016 | [22] |
| | Wurz "Auf der Schlattein" | 24.813 | [22] |
| *NE* | *Q1* | *970* | |
| | *Q2* | *4959* | |
| | *Q3* | *14826* | |
| SE | | | |
| | Sattenbeuren-Kieswerk | 876 | [24] |
| | Abri Wachtfels | 2.529 | [23] |
| | Abri Unter den Seewänden | 2.857 | [26] pers. comm. B. Gehlen |
| | "Gemeindebeunden" Bad Buchau-Kappel | 3.056 | [27] |
| | Geispel | 3.474 | [23] |
| | Lengnau-Chlini Ey | 3.781 | [23]; pers. comm. J. Affolter / B. Gehlen |
| | Fru, La (423) | 3.864 | [41]; pers. comm. J. Affolter / B. Gehlen |
| | Feuerbichl - Schwangau-Horn | 4.807 | [29,42]; pers. comm. B. Gehlen |
| | Lüscherzmoos | 8.297 | [23]; pers. comm. J. Affolter / B. Gehlen |
| | Cham-Grindel III | 9.026 | [23] |
| | Langrüti | 10.309 | [23] |
| | Altwasser-Höhle 1 | 10.943 | [23] |
| | Fürsteiner | 15.239 | [23] |
| | Cham-Grindel I | 18.836 | [23] |
| *SE* | *Q1* | *3160* | |
| | *Q2* | *4336* | |
| | *Q3* | *9988* | |

| Region | Site | RMCA in km$^2$ | Reference |
|---|---|---|---|
| SW | | | |
| | Champ Chalatras | 1.524 | [43] |
| | Borie del Rey (426), La | 1.982 | [44] |
| | Balzola, Cueva de | 2.401 | [32] |
| | Urratxa III | 2.479 | [45] |
| | Troubat, La Grotte-Abri de/ Moulin, La grott | 2.522 | [33] |
| | Cuze de sainte-Anastasie UA4 | 6.153 | [46] |
| | Peyrazet, Le Grotte Abri de | 7.172 | [47,48] |
| | Port-de-Penne | 15.068 | [44] |
| | Mas d'Azil | 40.446 | [36] |
| | Cuze de sainte-Anastasie UA5 | 49.828 | [46] |
| *SW* | *Q1* | *2420* | |
| | *Q2* | *4337* | |
| | *Q3* | *13094* | |

The table comprises only data that were considered for calculation in this study (n = 39). Below each region, the corresponding quartiles (Q1, Q2, Q3) used to calculate the palaeodemographic estimates are shown (see **table 2**).

**S3 Table. Regionally distinguished demographic estimates for GI-1d-a based on dataset B (i.e., only directly dated or specifically attributable to GI-1d-a).**

| Core Area region | ODI (km$^2$) | Q | RMCA (km$^2$) | $N_{raw}$ | $N_{groups}$ | $N_{persons}$ | $D_{population}$ |
|---|---|---|---|---|---|---|---|
| | | 1 | 2,670 | | 18.3 | 777 | 0.016 |
| Great Britain | 48,810 | 2 | 5,117 | 17 | **9.5** | **405** | **0.008** |
| | | 3 | 5,896 | | 8.3 | 352 | 0.007 |
| | | 1 | | | 10.3 | 436 | 0.016 |
| N France | 27,421 | 2 | | | **5.4** | **228** | **0.008** |
| | | 3 | | | 4.7 | 198 | 0.007 |
| | | 1 | | | 36.9 | 1,567 | 0.016 |
| Benelux & NW Germany | 98,467 | 2 | | | **19.2** | **818** | **0.008** |
| | | 3 | | | 16.7 | 710 | 0.007 |
| | | 1 | | | 7.3 | 308 | 0.016 |
| S Scandinavia | 19,372 | 2 | | | **3.8** | **161** | **0.008** |
| | | 3 | | | 3.3 | 140 | 0.007 |
| | | 1 | | | 34.7 | 1,473 | 0.016 |
| Poland & NE Germany | 92,553 | 2 | | | **18.1** | **769** | **0.008** |
| | | 3 | | | 15.7 | 667 | 0.007 |
| | | 1 | 1,057 | | 34.2 | 1,455 | 0.040 |
| Czech Rep. & SE Germany | 36,181 | 2 | 1,948 | 10 | **18.6** | **789** | **0.022** |
| | | 3 | 2,626 | | 13.8 | 585 | 0.016 |
| | | 1 | 2,604 | | 7.6 | 323 | 0.016 |
| Switzerland & SW Germany | 19,783 | 2 | 3,550 | 19 | **5.6** | **237** | **0.012** |
| | | 3 | 8,662 | | 2.3 | 97 | 0.005 |
| | | 1 | | | 5.3 | 224 | 0.016 |
| Italy | 13,745 | 2 | | | **3.9** | **165** | **0.012** |
| | | 3 | | | 1.6 | 67 | 0.005 |
| | | 1 | | | 7.9 | 334 | 0.016 |
| SE France | 20,470 | 2 | | | **5.8** | **245** | **0.012** |
| | | 3 | | | 2.4 | 100 | 0.005 |
| | | 1 | 1,578 | | 23.4 | 996 | 0.027 |
| SW France | 36,993 | 2 | 2,522 | 7 | **14.7** | **624** | **0.017** |
| | | 3 | 20,730 | | 1.8 | 76 | 0.002 |
| | | 1 | | | 41.6 | 1,768 | 0.027 |
| Spain & French Pyrenees | 65,661 | 2 | | | **26.0** | **1,107** | **0.017** |
| | | 3 | | | 3.2 | 135 | 0.002 |
| | | 1 | | | 6.1 | 261 | 0.027 |
| Portugal | 9,700 | 2 | | | **3.8** | **163** | **0.017** |
| | | 3 | | | 0.5 | 20 | 0.002 |
| | | 1 | | | ***234*** | ***9,924*** | ***0.020*** |
| **Sum Core Area** | ***489,156*** | 2 | | **53** | **134** | **5,710** | **0.012** |
| | | 3 | | | ***74*** | ***3,147*** | ***0.006*** |
| | | 1 | | | | | ***0.004*** |
| **Total Area of Calculation** | ***2,600,000*** | 2 | | | | | **0.002** |
| | | 3 | | | | | ***0.001*** |

Thin horizontal lines indicate regions with transfer of RMCA data (cf. **S3 Fig**). ODI = optimally describing Isoline in km$^2$; Q = Quartile [1 = lower, 2 = median, 3 = upper]; RMCA (km$^2$) = raw material catchment area in km²; $N_{raw}$ = number of RMCAs; $N_{groups}$ = number of groups; $N_{persons}$ = number of persons; $D_{population}$ = population density within Core Areas as p/km$^2$, and the Total Area of Calculation (bottom rows).

**S4 Table. Regionally distinguished demographic estimates for GS-1 based on dataset B (i.e., only directly dated or specifically attributable to GS-1).**

| Core Area region | ODI (km²) | Q | RMCA (km²) | $N_{raw}$ | $N_{groups}$ | $N_{persons}$ | $D_{population}$ |
|---|---|---|---|---|---|---|---|
| | | 1 | 2,934 | | *8.4* | *358* | *0.014* |
| Great Britain | 24,720 | 2 | 5,187 | 4 | **4.8** | **203** | **0.008** |
| | | 3 | 7,462 | | *3.3* | *141* | *0.006* |
| | | 1 | | | *1.1* | *49* | *0.014* |
| N France | 3,360 | 2 | | | **0.6** | **28** | **0.008** |
| | | 3 | | | *0.5* | *19* | *0.006* |
| | | 1 | | | *17.4* | *740* | *0.014* |
| Benelux & NW Germany | 51,060 | 2 | | | **9.8** | **418** | **0.008** |
| | | 3 | | | *6.8* | *291* | *0.006* |
| | | 1 | | | *0.4* | *18* | *0.014* |
| S Scandinavia | 1220 | 2 | | | **0.2** | **10** | **0.008** |
| | | 3 | | | *0.2* | *7* | *0.006* |
| | | 1 | | | *49.0* | *2,085* | *0.014* |
| Poland & NE Germany | 143,890 | 2 | | | **27.7** | **1,179** | **0.008** |
| | | 3 | | | *19.3* | *819* | *0.006* |
| | | 1 | 970 | | *10.1* | *429* | *0.044* |
| Czech Rep. & SE Germany | 9,800 | 2 | 4,959 | 11 | **2.0** | **84** | **0.009** |
| | | 3 | 14,826 | | *0.7* | *28* | *0.003* |
| | | 1 | 3,160 | | *8.5* | *361* | *0.013* |
| Switzerland & SW Germany | 26,880 | 2 | 4,336 | 14 | **6.2** | **263** | **0.010** |
| | | 3 | 9,988 | | *2.7* | *114* | *0.004* |
| | | 1 | | | *9,5* | *405* | *0,013* |
| Italy | 30,120 | 2 | | | **6,9** | **295** | **0,010** |
| | | 3 | | | *3,0* | *128* | *0,004* |
| | | 1 | | | *3,2* | *136* | *0,013* |
| SE France | 10090 | 2 | | | **2,3** | **99** | **0,010** |
| | | 3 | | | *1,0* | *43* | *0,004* |
| | | 1 | 2,420 | | *9,4* | *401* | *0,018* |
| SW France | 22,860 | 2 | 4,337 | 10 | **5,3** | **224** | **0,010** |
| | | 3 | 13,094 | | *1,7* | *74* | *0,003* |
| | | 1 | | | *25,0* | *1065* | *0,018* |
| Spain & French Pyrenees | 60,630 | 2 | | | **14,0** | **594** | **0,010** |
| | | 3 | | | *4,6* | *197* | *0,003* |
| | | 1 | | | *3.1* | *133* | *0.018* |
| Portugal | 7,560 | 2 | | | **1.7** | **74** | **0.010** |
| | | 3 | | | *0.6* | *25* | *0.003* |
| | | 1 | | | ***145*** | ***6179*** | ***0,016*** |
| **Sum Core Area** | ***392,190*** | 2 | | **39** | **82** | **3471** | **0,009** |
| | | 3 | | | ***44*** | ***1886*** | ***0,005*** |
| | | 1 | | | | | ***0.002*** |
| **Total Area of Calculation** | ***2,600,000*** | 2 | | | | | **0.001** |
| | | 3 | | | | | ***0.001*** |

Thin horizontal lines indicate regions with transfer of RMCA data (cf. **S4 Fig**). For explanation on abbreviations see **S2 Table**.

**S5 Table. Number of radiocarbon-dated sites divided by GI-1d-a and GS-1**.

| | **period** | **ky (duration)** | **n of radiocarbon dates** | **%** | **n normalised to 1.3 ky** | **%** |
|---|---|---|---|---|---|---|
| all radiocarbon dates | GS-1 | 1.2 | 522 | 30 | 566 | 31 |
| | GI-1d-a | 1.3 | 1236 | 70 | | 69 |
| **Sum** | | | **1758** | | | |

| **region** | **period** | **years (duration)** | **presence of radiocarbon dates per site and time-bin** | **% per region** | **normalised to 1.3 ky** | **% per region** |
|---|---|---|---|---|---|---|
| Total Area of Calculation | GS-1 | 1.2 | 276 | 38 | 302 | 40 |
| | GI-1d-a | 1.3 | 455 | 62 | | 60 |
| **Sum** | | | **731** | | | |
| Great Britain | GS-1 | | 29 | 45 | 31 | 47 |
| | GI-1d-a | | 36 | 55 | | 53 |
| N France >47.4°N | GS-1 | | 12 | 41 | 13 | 43 |
| | GI-1d-a | | 17 | 59 | | 57 |
| Benelux & W Germany | GS-1 | | 26 | 29 | 28 | 31 |
| | GI-1d-a | | 63 | 71 | | 69 |
| S Scandinavia | GS-1 | | 7 | 39 | 8 | 41 |
| | GI-1d-a | | 11 | 61 | | 59 |
| Poland & NE Germany | GS-1 | | 29 | 37 | 31 | 39 |
| | GI-1d-a | | 49 | 63 | | 61 |
| Czech Rep. & SE Germany | GS-1 | | 6 | 25 | 7 | 27 |
| | GI-1d-a | | 18 | 75 | | 73 |
| Switzerland | GS-1 | | 3 | 18 | 3 | 19 |
| | GI-1d-a | | 14 | 82 | | 81 |
| Italy | GS-1 | | 28 | 44 | 30 | 46 |
| | GI-1d-a | | 36 | 56 | | 54 |
| SE France >1.8°E | GS-1 | | 39 | 42 | 42 | 44 |
| | GI-1d-a | | 53 | 58 | | 56 |
| SW France <1.8°E | GS-1 | | 20 | 36 | 22 | 38 |
| | GI-1d-a | | 35 | 64 | | 62 |
| Spain & French Pyrenees | GS-1 | | 70 | 38 | 76 | 40 |
| | GI-1d-a | | 114 | 62 | | 60 |
| Portugal | GS-1 | | 7 | 44 | 8 | 46 |
| | GI-1d-a | | 9 | 56 | | 54 |

The numbers are given for the entire database (“all radiocarbon dates”, upper), for the Total Area of Calculation (middle) and for each region (lower). Dates were taken from the Radiocarbon Palaeolithic Europe Database v.27 [49], see details in main publication. Except for the first dataset (n = 1758), radiocarbon dates per site and time-bin were counted as one occurrence each (n = 731). To account for the slightly shorter phase of the GS-1 (1.2 ky) compared to GI-1d-a (1.3 ky), the GS-1 radiocarbon counts were normalised for the duration. Note that the regions slightly differ from the ones defined in **Fig 1** (that is for “Benelux and W Germany” and Switzerland).

### Figures

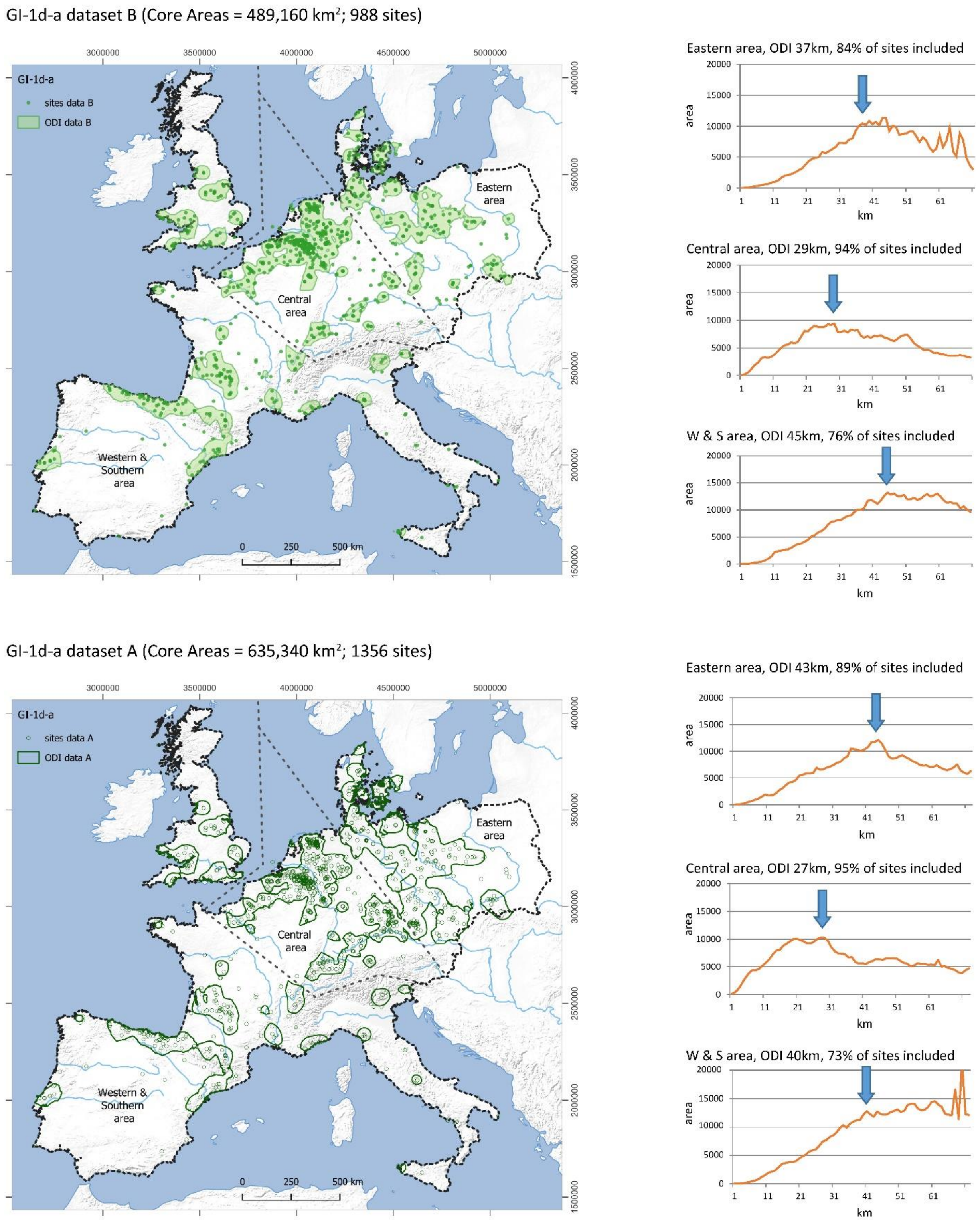


**S1 Fig. Maps (left) and diagrams (right) from the definition of the Core Areas based on the sites dated to Greenland Interstadial (GI) 1d-a.** The maps of dataset A (upper, sites n=1356) and dataset B (lower, n=988) include dashed lines which divide areas for which Core Areas (green areas) – areas of equal site densities – were calculated separately (see [1]), the dotted line delimits the Total Area of

Calculation. Regional areal increase of isolines (right) was interpolated by using distance values of Largest Empty Circles for GI-1d-a (right). X-axis of diagrams display radius of Largest Empty Circle, y-axis areal increase ($km^2$) encircled by the Isoline. To define the Optimal Describing Isoline (ODI, arrows in diagrams), the first peak or plateau, encircling around 70% or more of all sites, is selected. Core Areas of dataset A comprises 90% of the sites, Core Areas of dataset B 85%.

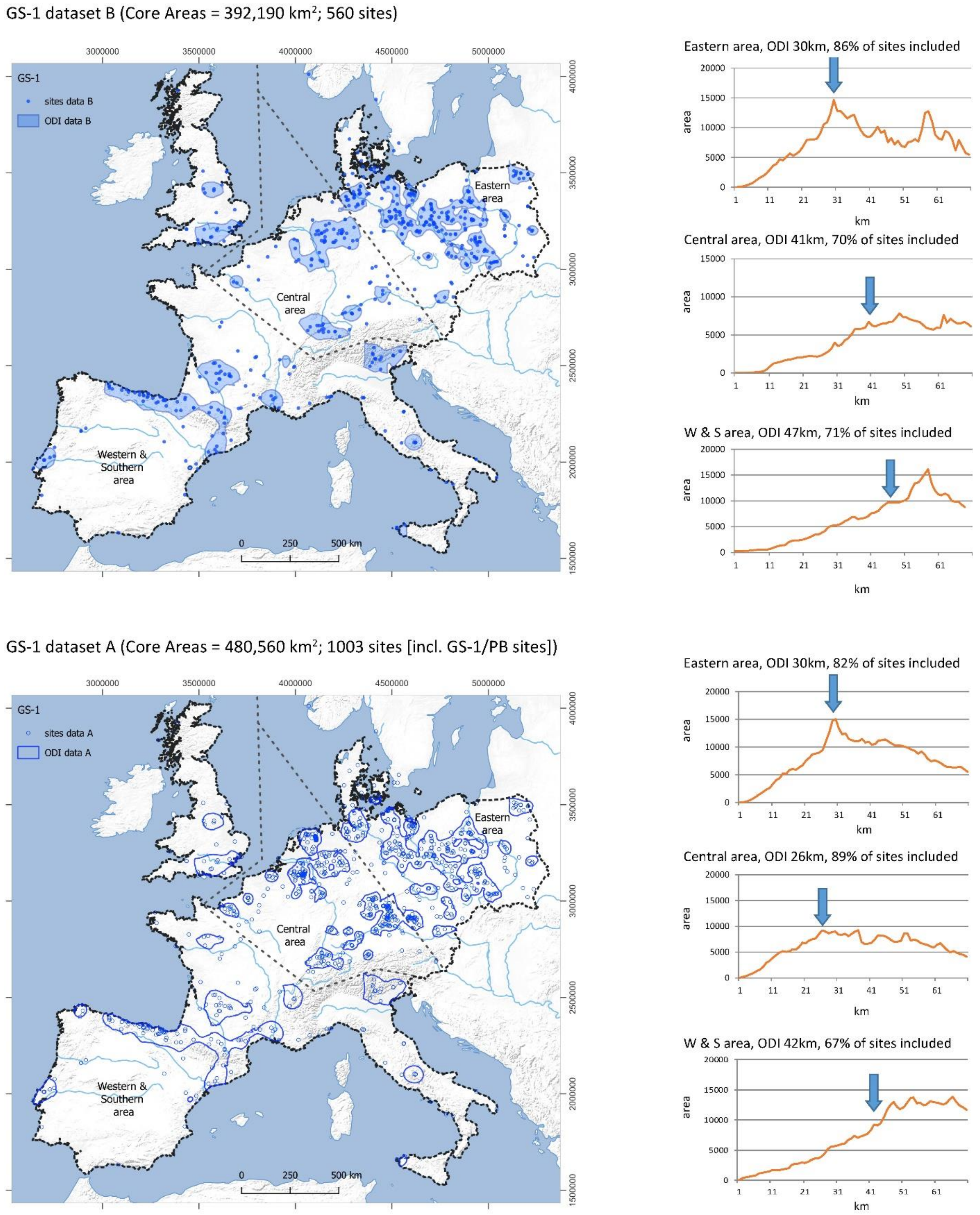


**S2 Fig. Maps (left) and diagrams (right) from the definition of the Core Areas based on the sites dated to Greenland Stadial (GS) 1.** The maps of dataset A (upper, sites n= 1003) and dataset B (lower, n=560) include dashed lines which divide areas for which Core Areas (blue areas) – areas of equal site densities – were calculated separately (see [1]). Dataset A comprises 82% of the sites, Core Areas of dataset B 74%. For explanation on details see **S1 Fig**.

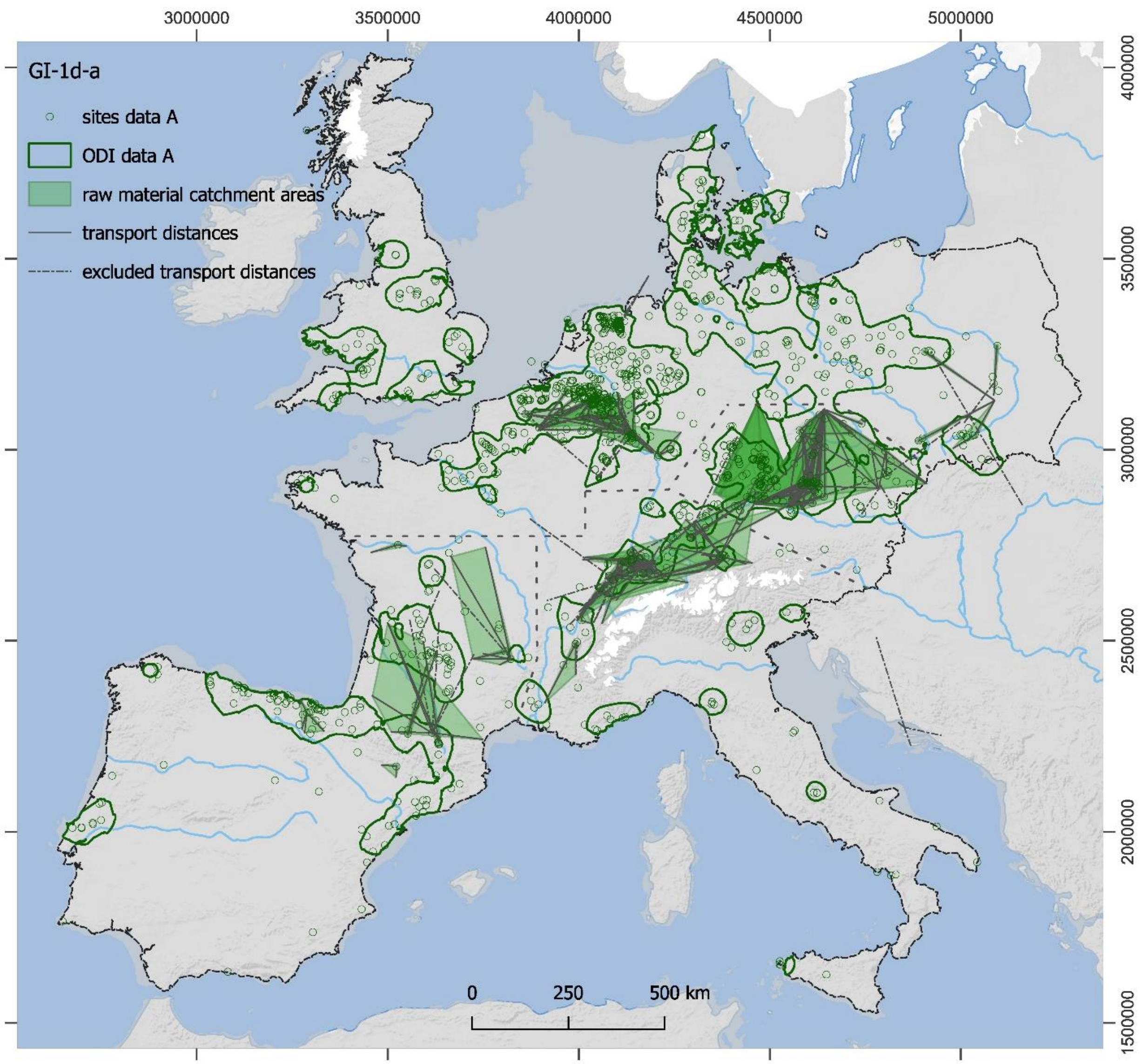


**S3 Fig. Map of all Raw Material Catchment Areas (RMCA) from sites dated to GI-1d-a that were collected from the literature (n = 108).** Several RMCAs were excluded from further analysis (see explanation in main article and remaining sites [n = 53] in **table S1**). Thick dashed lines delimit regions within which the same RMCA data was used to calculate the demographic estimates. Background: Core Areas and sites from dataset A.

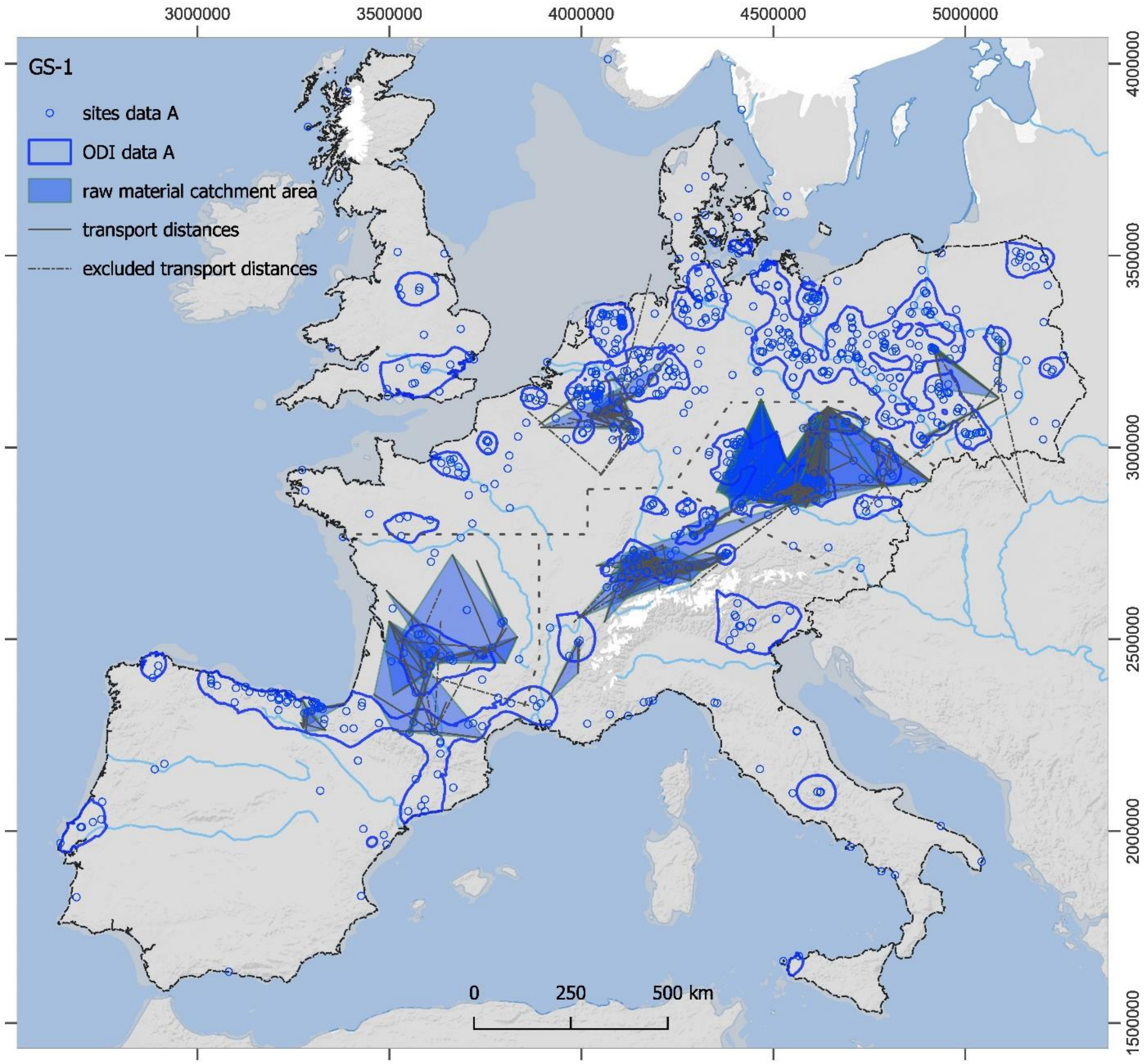


**S4 Fig. Map of all Raw Material Catchment Areas (RMCA) from sites dated to GS-1 that were collected from the literature (n = 97).** Several RMCAs were excluded from further analysis (see explanation in main article and remaining sites [n = 39] in **S2 Table**). Thick dashed lines delimit regions within which the same RMCA data was used to calculate the demographic estimates. Background: Core Areas and sites from dataset A.

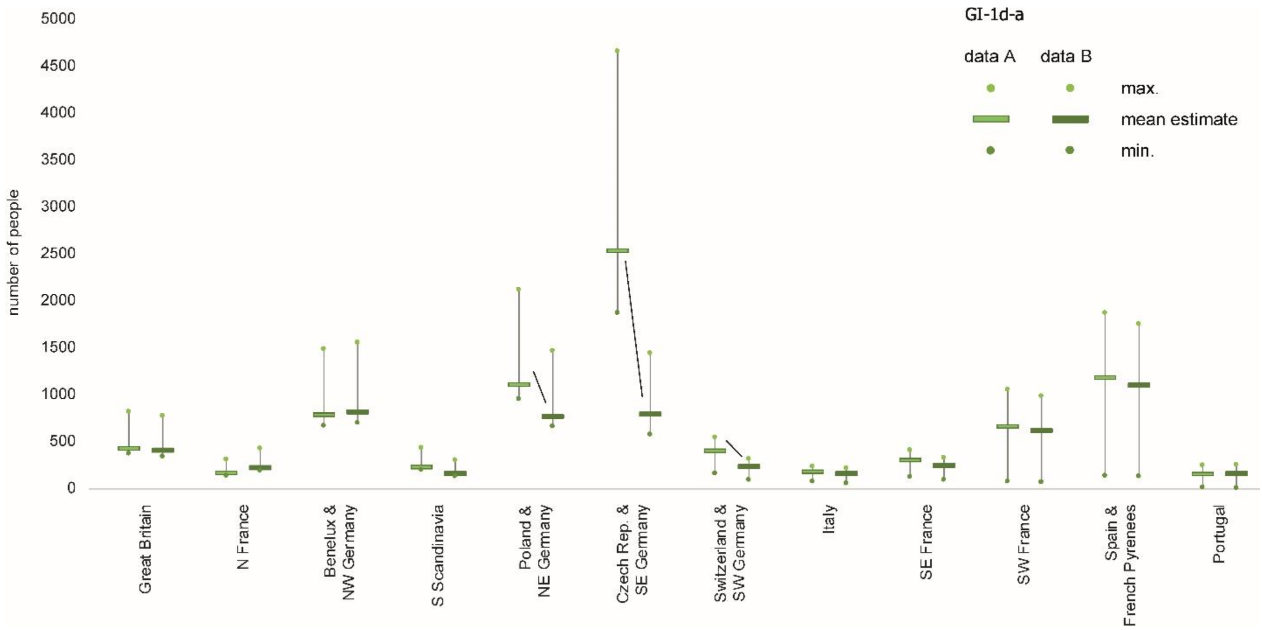


**S5 Fig. GI-1d-a: Comparison of estimates of people for dataset A (left bar) and B (right bar) per region.** The largest discrepancies occur in areas of NE, SE and SW Germany and adjacent regions.

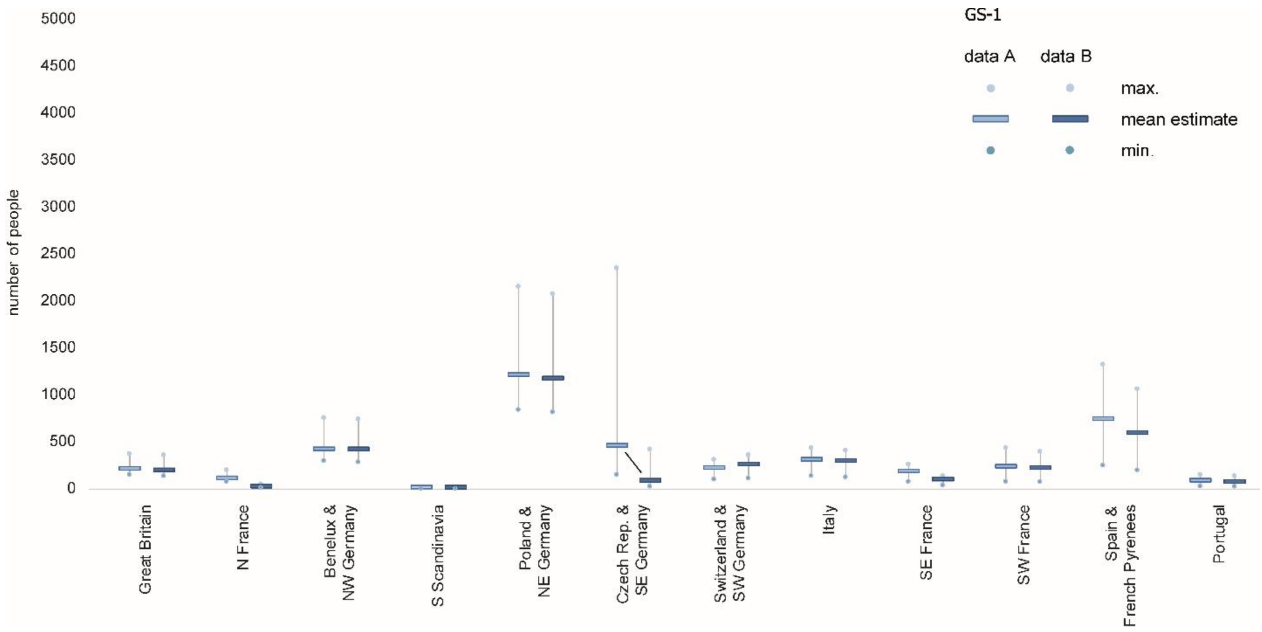


**S6 Fig. GS-1: Comparison of estimates of people for dataset A (left bar) and B (right bar) per region.** Discrepancies occur in areas the Czech Republic and SE Germany.

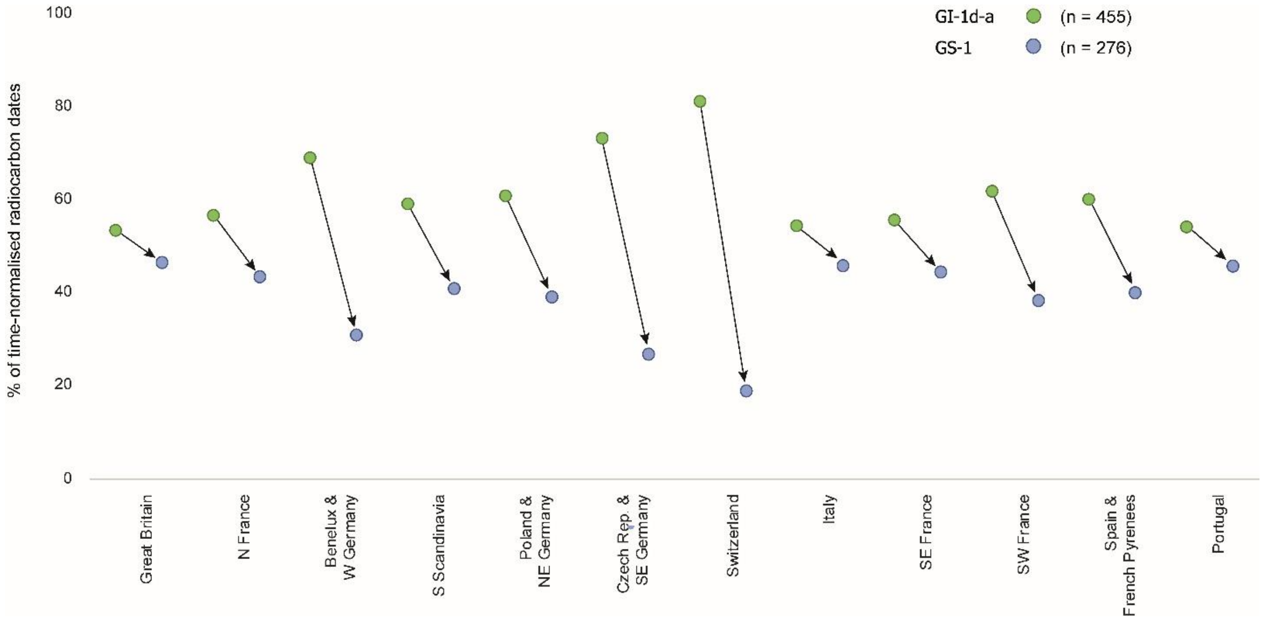


**S7 Fig**. **Diachronic tendencies in the frequency of regionally differentiated radiocarbon dates per region**. The percentage of dates for GI-1d-a (green dot, left side of the arrow) and GS-1 (blue dot, right) are displayed for each region. Dates were taken from the Radiocarbon Palaeolithic Europe Database v.27 [49]. We counted each time-bin covered by radiocarbon date(s) from an archaeological site as one occurrence (see **S3 Table**). To account for the slightly shorter phase of the GS-1 compared to GI-1d-a, the GS-1 radiocarbon counts were normalised for the duration of the phase.

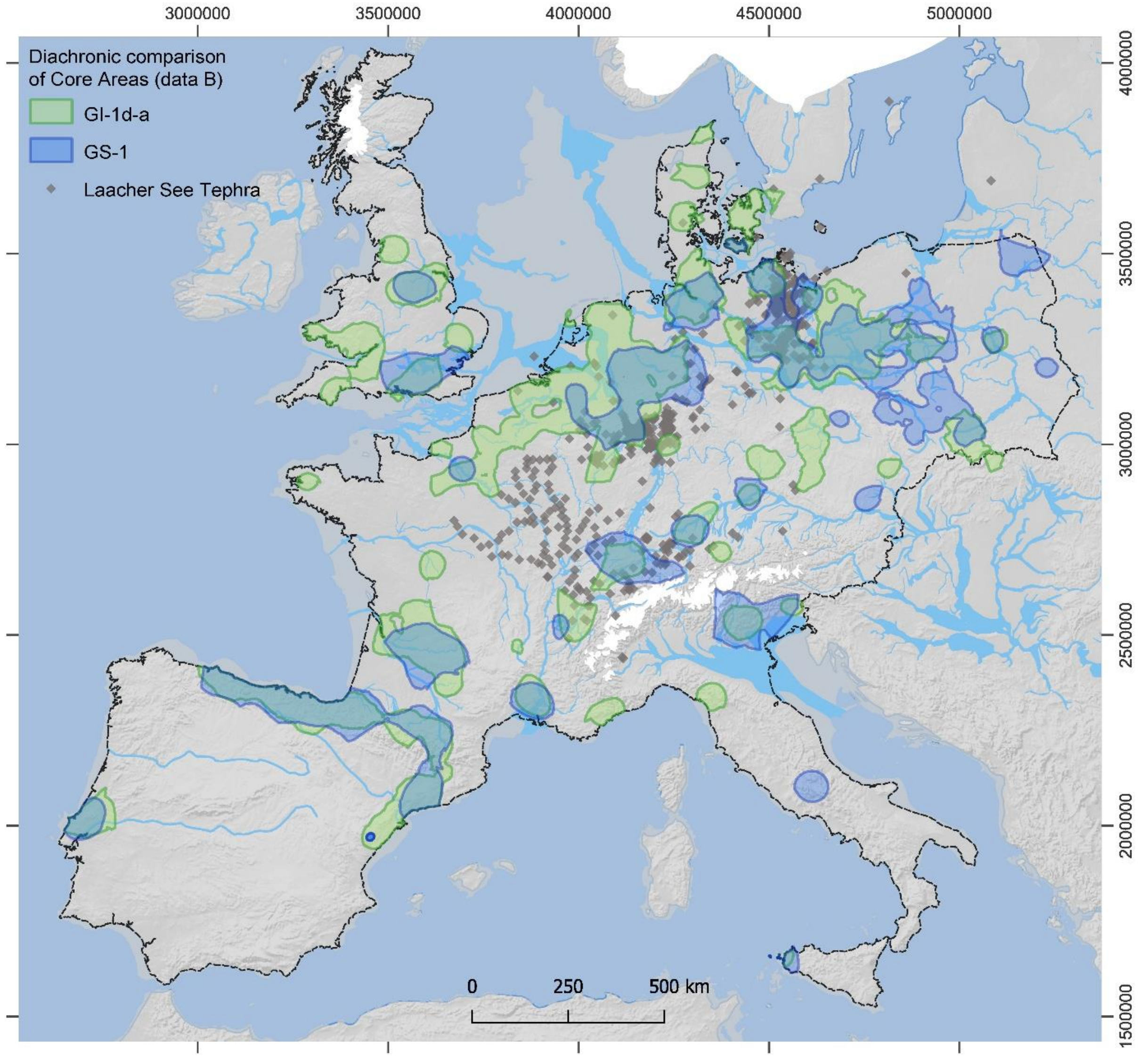


**S8 Fig. Comparison of Final Palaeolithic Core Areas (data B) from GI-1d-a (green areas) to GS-1 (blue areas).** For a comparison of data A Core Areas, see **Fig 7**. Grey diamonds indicate documented (micro)tephra from the Laacher See eruption, dated to 13 ky BP.

### Supporting information References

8. Heinen M. Der Federmesser-Horizont am Niederrhein und im angrenzenden Mittelgebirgsraum: Regionale und interne Organisation. In: Baales M, Pasda C, editors. „All der holden Hügel ist keiner mir fremd.": Festschrift zum 65 Geburtstag von Claus-Joachim Kind. Bonn: Rudolf Habelt; 2019.

9. Stoop D. Federmesser mobility patterns in the Western Meuse area, Limburg, the Netherlands: the case studies of Horn-Haelen and Heythuysen-de Fransman I. Leiden: Leiden University; 2014. Available: https://hdl.handle.net/1887/28507

10. Baales M. Archäologie des Eiszeitalters: frühe Menschen an Mittelrhein und Mosel. Koblenz: Gesellschaft für Archäologie an Mittelrhein und Mosel; 2005.

11. Baales M. Neue Untersuchungen zum Spätpaläolithikum des Neuwieder Beckens : einige Aspekte des Federmesser-Fundplatzes Kettig, Kr. Mayen-Koblenz. Den Bogen spannen: Festschrift für Bernhard Gramsch zum 65 Geburtstag. Weissbach: Beier & Beran; 1999. pp. 55–66.

12. Floss H. Rohmaterialversorgung im Paläolithikum des Mittelrheingebietes. Bonn: Habelt; 1994.

13. Loew S. Rüsselsheim 122 und die Federmessergruppen am Unteren Main. Ph.D., Universität zu Köln. 2006. Available: http://kups.ub.uni-koeln.de/id/eprint/2079

14. Crombé P, Sergant J, Verbrugge A, De Graeve A, Cherretté B, Mikkelsen J, et al. A sealed flint knapping site from the Younger Dryas in the Scheldt valley (Belgium): Bridging the gap in human occupation at the Pleistocene–Holocene transition in W Europe. J Archaeol Sci. 2014;50: 420–439. doi:10.1016/j.jas.2014.07.021

15. Baales M. Umwelt und Jagdökonomie der Ahrensburger Rentierjäger im Mittelgebirge. Bonn: Dr. Rudolf Habelt GmbH; 1996.

16. Sulgostowska Z. Final Palaeolithic Societies' Mobility in Poland as Seen from the Distribution of Flints. Arch Baltica. 2006;7: 36–42.

17. Oliva M. Encyklopedie paleolitu a mezolitu českých zemí. Vydání 1. Brno: Moravské zemské muzeum; 2016.

18. Tarbska J, Walanus A, Ciesielczuk J, Samek L, Dutkiewicz E. Ferruginous Raw Material Sources for Palaeolithic in Poland (Central Europe)? Provenance Studies: Occurrence, Litostratigraphy and Application. Jerusalem; 2008. Available: https://www.ndt.net/?id=6175

19. Vencl S. Prehistory of Bohemia 1: The Palaeolithic and Mesolithic. Praha: Archeologický ústav AV CR; 2013.

20. Stefański D, Wilczyński J. Extralocal raw materials in the Swiderian Culture: Case study of Kraków-Bieżanów sites. Anthropologie. 2012;50: 427–442.

21. Šída P, Pokorný P. Determining the archaeological potential of the landscape using Quaternary geological mapping in the Třebon region, south Bohemia. Arch Rozhledy. 2011;63: 485.

22. Sauer F. Late Palaeolithic Land Use Patterns in Bavaria. Ph.D., Friedrich-Alexander-Universität. 2018.

23. Affolter J. Silexrohstoffe - Schlüssel zu Analyse von Beziehungsnetzen. Die letzten Wildbeuter der Eiszeit - Neue Forschungen zum Spätpaläolithikum im Kanton Basel-Landschaft. Bern: Schwabe Verlag; 2015. pp. 198–209.

24. Kind C-J. Sattenbeuren - Kieswerk, ein spätpaläolithischer Uferrandlagerplatz am Federsee. Fundb Baden-Württemberg. 1995;20: 159–194.

25. Nielsen EH, Affolter J. Wauwil Station 25-Sandmatt: eine spätpaläolithische Fundstelle im Wauwilermoos. Luzern: Kantonaler Lehrmittelverlag; 1999.

35. Surmely F, Quinqueton A, Virmont J. Le gisement épipaléolithique ancien de la grotte Béraud à Saint-Privat-d'Allier (Haute-Loire, France). hal-00350923 , version 1. 2001.

36. Kegler JF. Das Azilien von Mas d'Azil. Der chronologische und kulturelle Kontext der Rückenspitzengruppen in Südwesteuropa. Mit einem Beitrag von Jan F. Kegler und Stefan R. Loew. Ph.D., Universität zu Köln. 2007. Available: http://kups.ub.uni-koeln.de/id/eprint/4231

37. Blank R. Ein Fundplatz der endpaläolithischen Stielspitzen - Gruppe am nördlichen Mittelgebirgsrand. Arch Korr. 1985;15: 287–292.

38. Balthasar P. Untersuchung des Wandels der Steinartefaktgrundproduktion in der Westfälischen Bucht vom Spätpaläolithikum bis zum Mesolithikum. Ph.D., Friedrich-Schiller-Universität. 2019.
